# Genetic instability from a single S-phase after whole genome duplication

**DOI:** 10.1101/2021.07.16.452672

**Authors:** Simon Gemble, René Wardenaar, Kristina Keuper, Nishit Srivastava, Maddalena Nano, Anne-Sophie Macé, Andréa E. Tijhuis, Sara Vanessa Bernhard, Diana C.J. Spierings, Anthony Simon, Oumou Goundiam, Helfrid Hochegger, Matthieu Piel, Floris Foijer, Zuzana Storchová, Renata Basto

**Author notes:** These authors contributed equally (René Wardenaar, Kristina Keuper).

## Abstract

Diploid and stable karyotypes are associated with health and fitness in animals. In contrast, whole genome duplications (WGDs) - doubling full chromosome content - are linked to genetic instability (GIN) and frequently found in human cancers ^1–3^. It has been established that WGDs fuel chromosome instability through abnormal mitosis ^4–8^, however, the immediate consequences of tetraploidy in the first interphase are not known. This is an essential question because single WGD events such as cytokinesis failure can promote tumorigenesis ^9^. Here, we found that newly born tetraploid human cells undergo high rates of DNA damage during DNA replication in the first S-phase. Using DNA combing and single cell sequencing, we show that DNA replication dynamics is perturbed, generating under- and over-replicated regions. Mechanistically, we found that these defects result from the lack of protein mass scaling up at the G1/S transition, which impairs the fidelity of DNA replication. This work shows that within a single interphase, unscheduled tetraploid cells can acquire highly abnormal karyotypes. These findings provide an explanation for the GIN landscape that favors tumorigenesis after tetraploidization.

---

Since WGDs can have different origins ^10, 11^, we developed several approaches to induce tetraploidization through either mitotic slippage (MS), cytokinesis failure (CF) or endoreplication (EnR) in the human diploid and genetically stable RPE-1 cell line. Most cells resulting from CF contained two nuclei, while EnR or MS generated mononucleated tetraploid cells. Cell size, number and nuclear size and centrosome number were considered, to distinguish diploids from tetraploids (Fig. 1a-b and Extended data Fig. 1a-i). For each approach, a mix of diploid and tetraploid cells was obtained, allowing the comparison of internal diploid controls and tetraploids. In all conditions, most tetraploid cells continued to cycle throughout the first interphase, allowing us to probe the consequences of tetraploidy within the first cell cycle.

**Figure 1:**
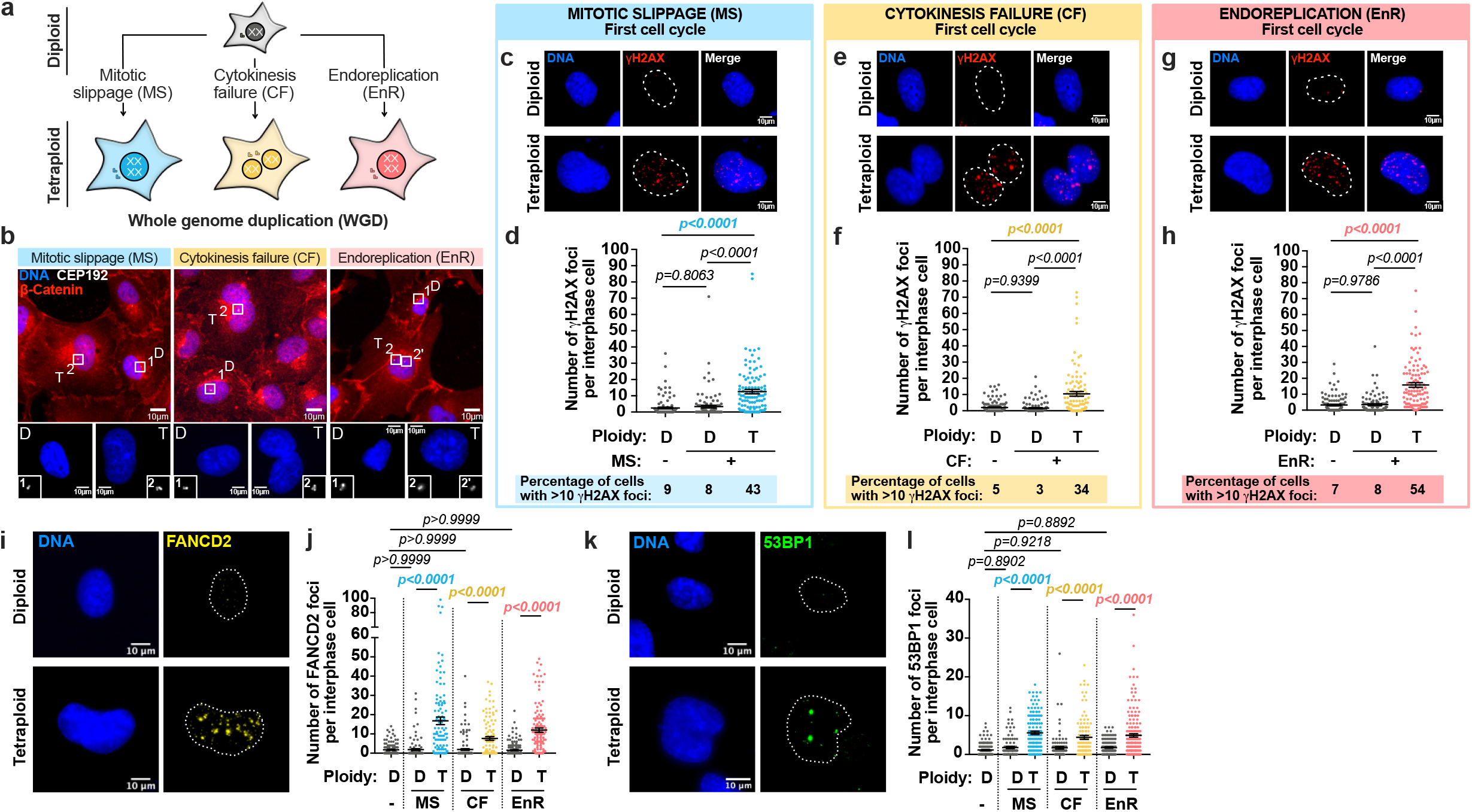
High levels of DNA damage in the first interphase following unscheduled WGD. **(a)** Schematic representation of the different means employed to generate tetraploid cells. **(b)** Images of diploid (D) and tetraploid (T) RPE-1 cells generated by mitotic slippage (MS), cytokinesis failure (CF) or endoreplication (EnR). DNA in blue, centrosomes labeled with anti-CEP192 antibodies (in white) and cell membranes with anti-β-Catenin antibodies (in red). The white squares correspond to the higher magnifications shown below. **(c, e** and **g)** Images showing DNA damage revealed by anti-γH2AX antibodies (in red) in diploid and tetraploid RPE-1 cells as indicated. DNA in blue. **(d, f** and **h)** Graphs showing the number of γH2AX foci per interphase cell in diploid and tetraploid RPE-1 cells. Mean ± SEM, >100 interphase cells, three independent experiments. The percentage of interphase cells with ≥10 γH2AX foci for each condition is indicated under the graph. **(i** and **k)** Images of diploid and tetraploid RPE-1 cells generated by MS labeled with anti-FANCD2 **(i)** or anti-53BP1 **(k)** antibodies (in yellow and green, respectively). DNA in blue. **(j** and **l)** Graphs showing the number of FANCD2 **(j)** or 53BP1 **(l)** foci per interphase cell in diploid and tetraploid RPE-1 cells. Mean ± SEM, >100 interphase cells, three independent experiments. The dotted lines indicate the nuclear area. **(d, f, h, j** and **l)** ANOVA test (one-sided).

Using an early marker of DNA damage - γH2AX - we found high levels of DNA damage in tetraploid cells (but not in controls) independently of the way they were generated (Fig. 1c-h, and Extended data Fig. 1a-i and Extended data Fig. 2a-f; methods). Moreover, while most diploid cells exhibited a low number of γH2AX foci (5-9%), the percentage of tetraploid cells with more than 10 foci was high (34-54%; Fig. 1c-h). The number of γH2AX foci correlated with fluorescence intensity (FI) levels (Extended data Fig. 1j). We excluded the possibility that the increase in tetraploid cells was simply due to increased nuclear size by normalizing the number of γH2AX foci by the nuclear area or nuclear FI (Extended data Fig. 1k-l). High levels of DNA damage were also found in tetraploid BJ fibroblast and HCT116 cells upon WGD (Extended data Fig. 2g-h).

To evaluate DNA damage levels after WGD, we compared DNA damage between tetraploid and diploid cells with replication stress (RS). RS results from the slowing or stalling of replication forks, which can be induced by high doses of Aphidicolin (APH), a DNA polymerase inhibitor, or Hydroxyurea (HU), a ribonucleotide reductase (RNR) inhibitor ^12, 13^. APH or HU generated comparable levels of DNA damage in diploid cells, when compared to untreated tetraploid cells (Extended data Fig. 2i). In addition to γH2AX, we also observed a significant increase in the number of FANCD2 and 53PB1 foci, which are double strand break repair factors ^14^, within the first interphase upon WGD (Fig. 1i-l). Further, tetraploid cells showed an increased olive tail moment in alkaline comet assays, revealing single and double strand breaks (Extended data Fig. 2j-k).

We next asked whether DNA damage is also generated in the subsequent cell cycles. A high proportion of tetraploid RPE-1 cells arrests after the first cell cycle in a LATS2-p53 dependent manner ^15^. We thus analyzed DNA damage levels in p53-depleted cells (Extended Fig. 2l). During the 2^nd^ and 3^rd^ interphases that followed tetraploidization, we observed a considerable decrease in DNA damage levels (Extended Fig. 2m-o). Since most animal cells are normally organized in tissues with cell-cell adhesions, we tested the consequences of WGDs in 3D cultures. Strikingly, 3D spheroids containing tetraploid cells displayed a higher γH2AX index (methods) compared to diploid cells (Extended data Fig 3a-d).

Collectively, our results show that a transition from a diploid to a tetraploid status after unscheduled WGD is accompanied by high levels of DNA damage within the first cell cycle.

## DNA replication-dependent DNA damage

We determined the cell cycle stage in which the DNA damage occurs using the fluorescence ubiquitination cell cycle indicator (FUCCI). During G1, the number of γH2AX foci was quite low and comparable to controls. As tetraploid cells enter S-phase, a slight increase in the number of foci could be observed, which increased substantially at the end of S-phase (Fig. 2a-b and Extended data Fig. 3e-f). These results were further confirmed by time lapse imaging using H2B-GFP to visualize DNA and 53BP1-RFP (Extended data Fig. 3g-h and Extended data Videos 1-2). To confirm that DNA damage in tetraploid cells appeared during S-phase, we blocked cells at the G1/S transition using high doses of either CDK4/6 or CDK2 inhibitors for 16hrs (Extended data Fig. 3i-j). We chose the 16hrs time period because it corresponds to the end of S-phase in the cycling population (Fig. 2a-b) and so it allows us to distinguish if DNA damage accumulates in a specific cell cycle phase or, alternatively after a certain period of time. G1-arrested tetraploid cells showed low levels of DNA damage, whereas cells released in S-phase exhibited high levels of DNA damage (Extended data Fig. 3i-o). Importantly, we observed a significant increase in the percentage of γH2AX foci co-localizing with markers of active DNA replication sites visualized by Proliferating Cell Nuclear Antigen (PCNA) and EdU incorporation in tetraploid cells compare to diploid cells (31% vs 7%; Extended data Fig. 3p-q).

**Figure 2:**
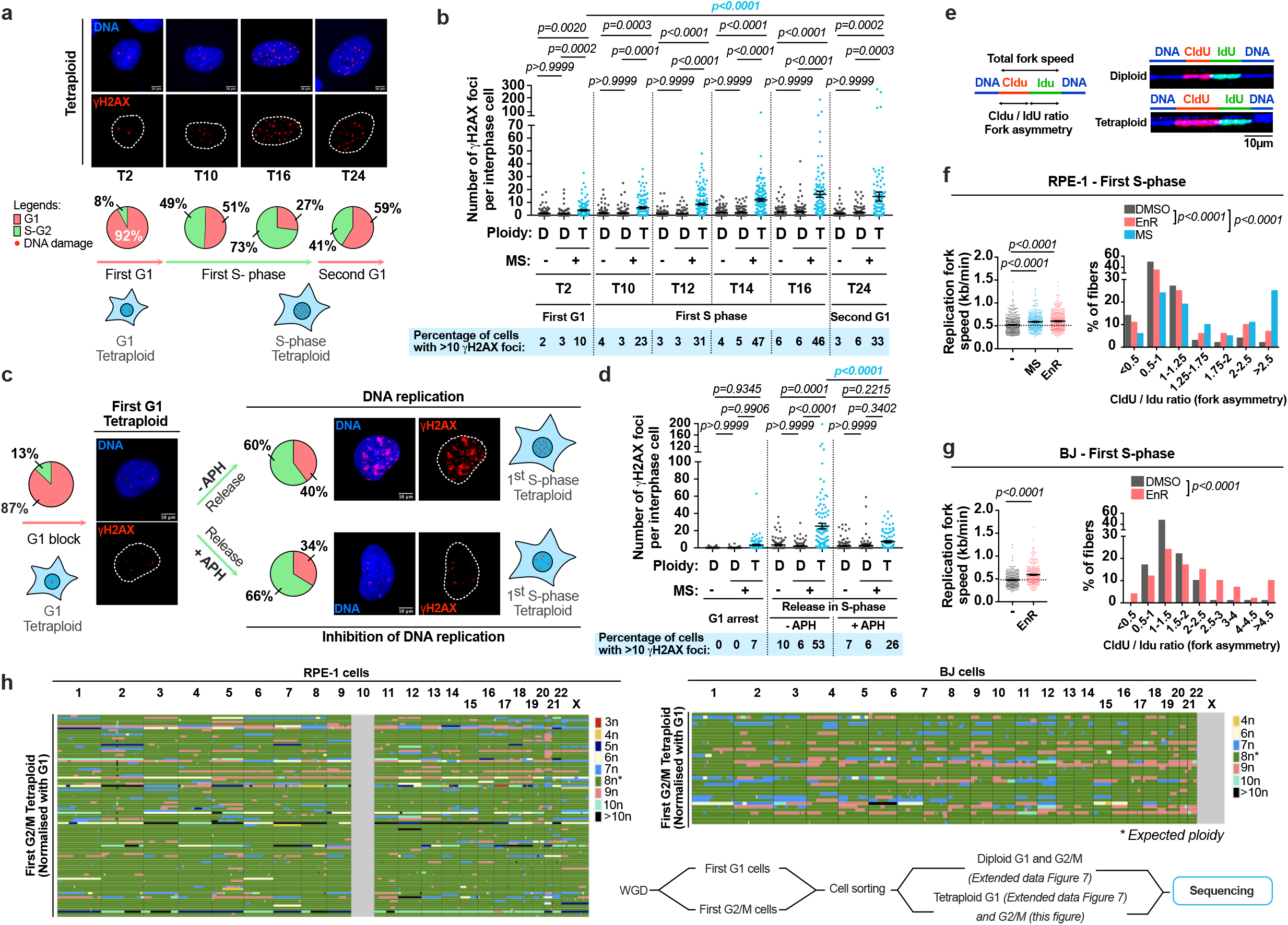
Genetic instability in tetraploid cells is generated during S-phase in a DNA replication-dependent manner. **(a)** Upper - Images showing DNA damage (anti-γH2AX antibodies in red) in RPE-1 tetraploid cells. DNA in blue. Lower -Percentage of RPE-1 tetraploid cells in G1 or in S-G2. Mean ± SEM, >100 interphase cells, three independent experiments. **(b)** Graph showing the number of γH2AX foci per interphase cell in diploid (D) and tetraploid (T) RPE-1 cells. Mean ± SEM, >100 interphase cells, three independent experiments. **(c)** Percentage of RPE-1 tetraploid cells in G1 or in S-G2 and representative images showing DNA damage (anti-γH2AX antibodies, red) in tetraploid cells synchronized in G1 using 1μM palbociclib or released in S-phase ± 400nM aphidicolin (APH). DNA in blue. Mean ± SEM, >100 interphase cells, three independent experiments. **(d)** Graph showing the number of γH2AX foci per interphase cell in diploid and tetraploid RPE-1 cells from **(c)**. Mean ± SEM, >100 interphase cells, three independent experiments. **(e)** Left - Scheme for replication fork analysis. Right - Immunofluorescence of DNA fibers obtained from diploid and tetraploid RPE-1 cells. **(f-g)** Left - Graph representing the replication fork speed in diploid and tetraploid RPE-1 (**f**) or BJ cells (**g**). Right - Graph showing the CldU/IdU ratio in diploid and tetraploid RPE-1 (**f**) or BJ cells (**g**). Mean ± SEM, **(f)** > 330 replication forks, **(g)** > 295 replication forks. **(i)** Genome-wide copy number plots for G2/M tetraploid RPE-1 or BJ cells induced through MS. Each row represents a cell. The copy number state is indicated in colors. Lower right - Schematic workflow showing the method used to sort the cells. **(b,d** and **f)** ANOVA test (one-sided). **(g)** t-test (two-sided).

By evaluating markers of DNA damage signaling and repair pathways we noticed that the number of KU80 and XRCC1 foci, proteins involved in Non-Homologous End Joining (NHEJ) ^16^ remained low in tetraploid cells. In contrast, the number of RAD51 foci, a homologous recombination (HR) factor, was increased. Moreover, the percentage of RAD51 foci co-localizing with γH2AX foci was significantly increased in tetraploid cells compare to diploid cells (14% vs 3%). The RS markers, Replication protein A (RPA) and FANCD2 were also increased and a significant increase in the percentage of colocalization with γH2AX foci was observed in tetraploid cells compare to diploid cells (40% vs 14%; Extended data Fig. 4a-k). Together, these results demonstrate that tetraploid cells experience high levels of DNA damage during S-phase, showing *bona fide* DNA damage and HR markers.

We hypothesized that DNA damage in tetraploid cells arises from errors occurring during DNA replication. To test this possibility, cells were arrested in G1 (Extended data Fig. 3k). We then released them in the presence of very low doses of APH or PHA, a Cdc7 inhibitor to inhibit DNA replication (detected by absence of EdU) without generating DNA damage (methods). This leads to DNA replication inhibition, albeit maintaining the biochemical activity typical of the S-phase nucleus. Strikingly, DNA damage levels were dramatically decreased in tetraploid cells (Figure 2c-d and Extended data Fig. 5a-f). Importantly, in the few tetraploid cells that escaped DNA replication inhibition - revealed by high EdU incorporation - a high number of γH2AX foci were still noticed (Extended data Fig. 5g-h). Together, these results establish that WGD generates DNA replication-dependent DNA damage. Deoxyribonucleoside triphosphate (dNTP) exhaustion leads to RS and GIN ^17^. We tested if supplying nucleosides rescued DNA damage defects described above. This was however not the case in cells or in an *in vivo* model of polyploidy generation (Extended data Fig. 5i-j and Extended data Fig. 10g). These results suggest that unscheduled WGD do not induce exhaustion of nucleoside levels as described in other oncogenic conditions ^17^.

We characterized DNA replication using RPE-1 cell lines stably expressing PCNA chromobodies (Supplemental information and methods). Quantitative 4D live imaging of DNA replication in diploid and tetraploid cells revealed striking decrease in the total number of PCNA foci and their volume, also confirmed with EdU foci (Extended data Fig. 6a-f). This suggests a lack of scaling up with DNA content and fewer active replication sites in tetraploid cells. Time lapse analysis of PCNA and FI was used as a readout of early and late S-phase ^18^, revealing a longer early S-phase period in tetraploid cells (Supplemental information, Extended data Fig. 6g-i and Extended data Videos 3-4). We next performed DNA combing, which allows the visualization of replication fork behavior in single DNA fibers ^13, 19^. Median fork speed and fork asymmetry (a readout of stalled or collapse forks) were increased in tetraploid cells (Fig. 2e-g and Extended data Fig. 6j-k). We attempted to analyze inter origin distance (IOD), as it is known that the number of active regions can influence fork speed ^20^. Although not significative, a trend for increased IOD could be noticed in tetraploid cells (possible explanation in methods).

To assess the type of karyotype generated in a single S-phase after WGD, we used single cell DNA sequencing (scDNAseq) (Supplementary information, Supplementary methods 1 and methods). Strikingly, in G2/M tetraploid cells over-duplicated chromosomes (> 10) were identified in addition to frequent over- and under-replicated regions (9n, 7n and 4n) (Fig. 2h and Extended data Fig. 7a-b). Both aneuploidy/heterogeneity scores and the proportion of the genome affected by aneuploidies were increased in G2/M tetraploid cells (Fig. 2h and Extended data Fig. 7a-d and methods). Our data establish that WGD generates abnormal karyotypes within a single S-phase.

## Non-optimal S-phase in tetraploid cells

It is expected that tetraploid cells scale up RNAs and protein content by a factor of 2. Using pyronin Y staining and quantitative phase imaging to measure total RNAs and protein content, we found a lack of scaling up in newly born tetraploid cells (Fig 3a-b and Extended Data Fig. 8a-c). We next tested the levels of key DNA replication factors. We developed protocols to sort tetraploids from diploids based on FUCCI and DNA content from a common cell population (Fig. 3c and Extended data Fig. 8d-e, methods). The same number of cells was loaded for diploid and tetraploid conditions and total protein extracts were probed by western blot. The chromatin associated H2B variant, the cytoskeleton component Actin and the membrane component β-Catenin showed an increase in the levels of these three proteins consistent with tetraploidization. In stark contrast, by using H2B as a readout of DNA content, we showed that essential G1 and S-phase DNA replication factors did not scale up in tetraploid cells (Fig. 3d-f and Extended data Fig. 8f-g). In addition, we probed the chromatin-bound fractions from the two cell populations. We tested origin recognition complex 1 (ORC1) ^21^, the minichromosome maintenance 2 (MCM2) helicase ^22^, Cdc10-dependent transcript 1 protein (CDT1) ^23^ and Cell division cycle 6 (CDC6) ^24^. These proteins are key members of pre-replication complexes (RCs) and are normally loaded in G1 during origin licensing. We also tested PCNA, CDC45 ^25^ and Treslin ^26^, which are required for the initiation of DNA replication. We further probed the levels of E2F1, a transcription factor which activates the expression of S-phase genes ^27–29^. With the exception of Treslin, the total levels of all these proteins did not show the expected scaling up in tetraploid cells (Fig. 3e-f). Concerning the chromatin fractions, pre-RCs, Treslin and CDC45 levels also did not scale up (Fig. 3g-h and Extended data Fig. 8h).

**Figure 3:**
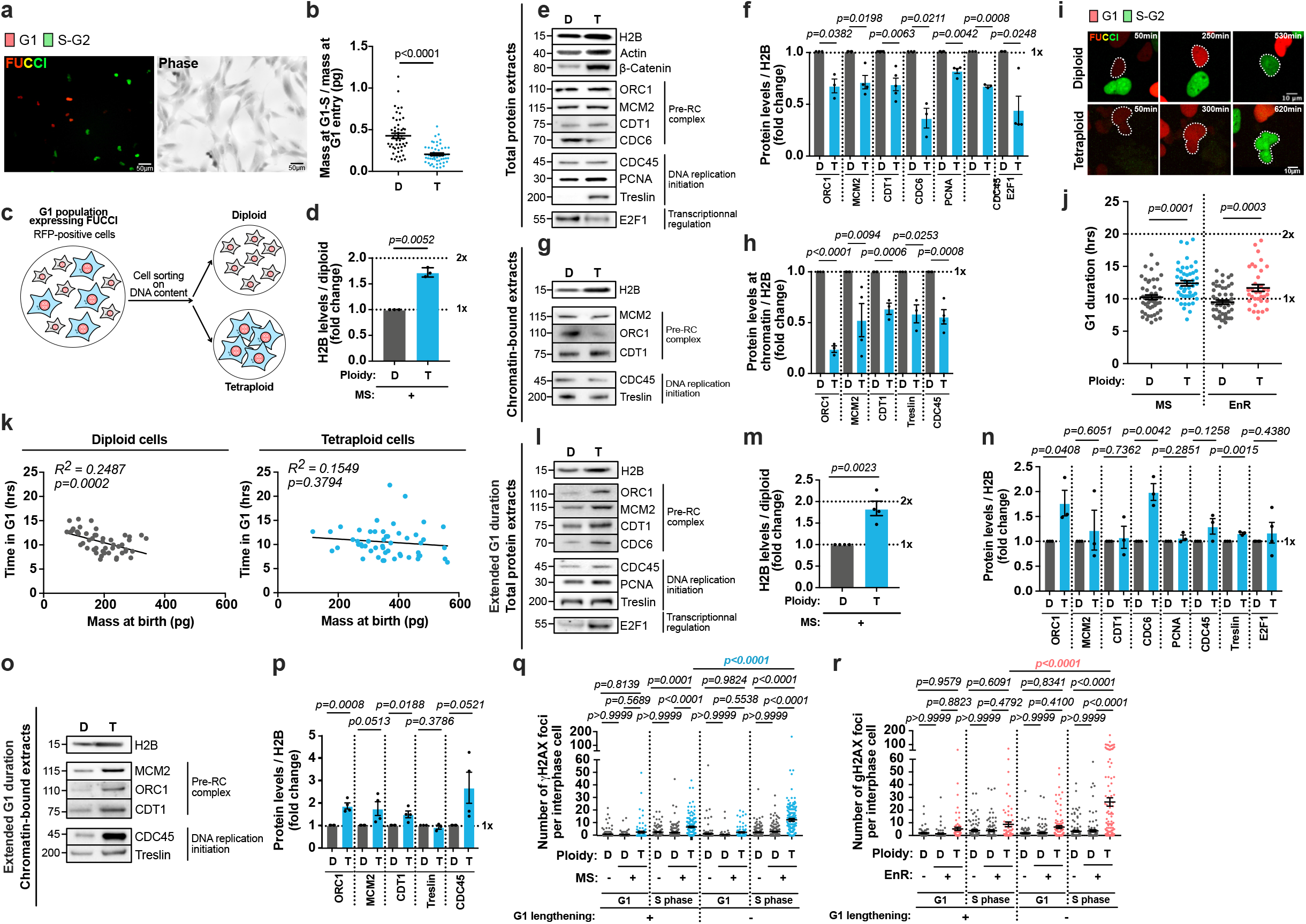
Key replication factors do not scale up in tetraploid cells. **(a)** Pictures of tetraploid cells expressing FUCCI and the correspondent signal in phase microscopy. **(b)** Graph representing the ratio of protein produced during G1 in diploid (D) and tetraploid (T) cells. Mean ± SEM, > 50 G1 cells, two experiments. **(c)** Schematic representation of FACS sorting. **(d)** Graph showing relative H2B levels in RPE-1 cells. Mean ± SEM, three experiments. **(e** and **g)** Immunoblots of total protein extracts **(e)** or chromatin bound extracts **(g)** obtained from RPE-1 cells. **(f** and **h)** Graph representing the protein levels from **(e** and **g)**. Mean ± SEM, three independent experiments. **(i)** Stills of time lapse Videos of RPE-1 cells expressing FUCCI. **(j)** Graph showing the G1 duration in RPE-1 cells. Mean ± SEM, > 35 interphase cells, two independent experiments. **(k)** Graphs showing the time in G1 and the mass at birth in RPE-1 cells. >50 interphase cells, two independent experiments. **(l** and **o)** Western blots of total protein extracts **(l)** or chromatin bound extracts **(o)** obtained from RPE-1 cells with extended G1 duration. **(m)** Graph showing relative H2B levels in RPE-1 cells with extended G1 duration. Mean ± SEM, four experiments. **(n** and **p)** Graphs representing the protein from **(l** and **o)**. Mean ± SEM, three experiments. **(q** and **r)** Graphs showing the number of γH2AX foci in RPE-1 cells with G1 lengthening or G1 arrest using 160nM/1μM palbociclib and released in S-phase. Mean ± SEM, >100 interphase cells, three experiments. **(e, g, l, o)**. The same number of cells was loaded for each condition. (**j, q** and **r**) ANOVA test (one-sided); (**d, f, h, m, n** and **o**) t-test (two-sided); (**k**) Pearson test (two-sided).

In normal proliferative cell cycles, growth occurring during G1 phase prepares cells for DNA replication allowing the expression and accumulation of key S-phase regulators ^28, 30^. We measured G1 duration in tetraploid cells to find only a slight increase when compared to diploid cells (Fig. 3i-j and Extended data Fig. 8i-j). Further, while a significant correlation between cell mass and G1 duration in diploid cells was detected, as described previously ^31^, this was absent in tetraploid cells (Fig. 3k). We then tested if G1 lengthening favored error-free DNA replication in tetraploid cells. We delayed S-phase entry using very low doses of CDK4/6 or CDK2 inhibitors (Extended data Fig. 9a-c and Supplemental information and methods). In this condition, the levels of DNA replication factors from total cell or chromatin extracts scaled up with DNA content (Fig. 3l-p vs 3e-h and Extended data Fig. 9k). Further, the number and volume of active replication sites in S-phase scaled up with DNA content in tetraploid cells and the dynamic behavior of PCNA in tetraploid cells was comparable to diploid cells (Extended data Fig. 9d-h and Extended data Videos 5-6). The time spent in S-phase was not altered but the ratio between early and late S-phase in tetraploid cells was restored (Extended data Fig. 9i-j). In all cell lines, G1 lengthening was sufficient to reduce the number of γH2AX, FANCD2 and 53BP1 foci in tetraploid S-phase cells (Fig. 3q-r and Extended data Fig. 9l-r).

Our data show that tetraploid cells transition from G1 to S-phase prematurely without undergoing scaling of global protein mass. They enter S-phase with insufficient DNA replication factors, which can be compensated for by G1 lengthening.

## E2F rescues GIN in tetraploid cells

Since the time spent in G1 does not prepare tetraploid cells for S-phase, we reasoned that increased E2F1 levels might compensate for defects in G1 length scaling up. E2F1 is a transcription factor that promotes proliferation and cell cycle progression by regulating S-phase and DNA replication factors ^28, 29^. We over-expressed E2F1 (OE) in diploid cells allowing us to increase the expression of DNA replication proteins just before generating tetraploid cells. Strikingly, this was sufficient to rescue the levels of DNA damage in tetraploid cells (Fig. 4a-c and Extended Data Fig. 10a-c).

**Figure 4:**
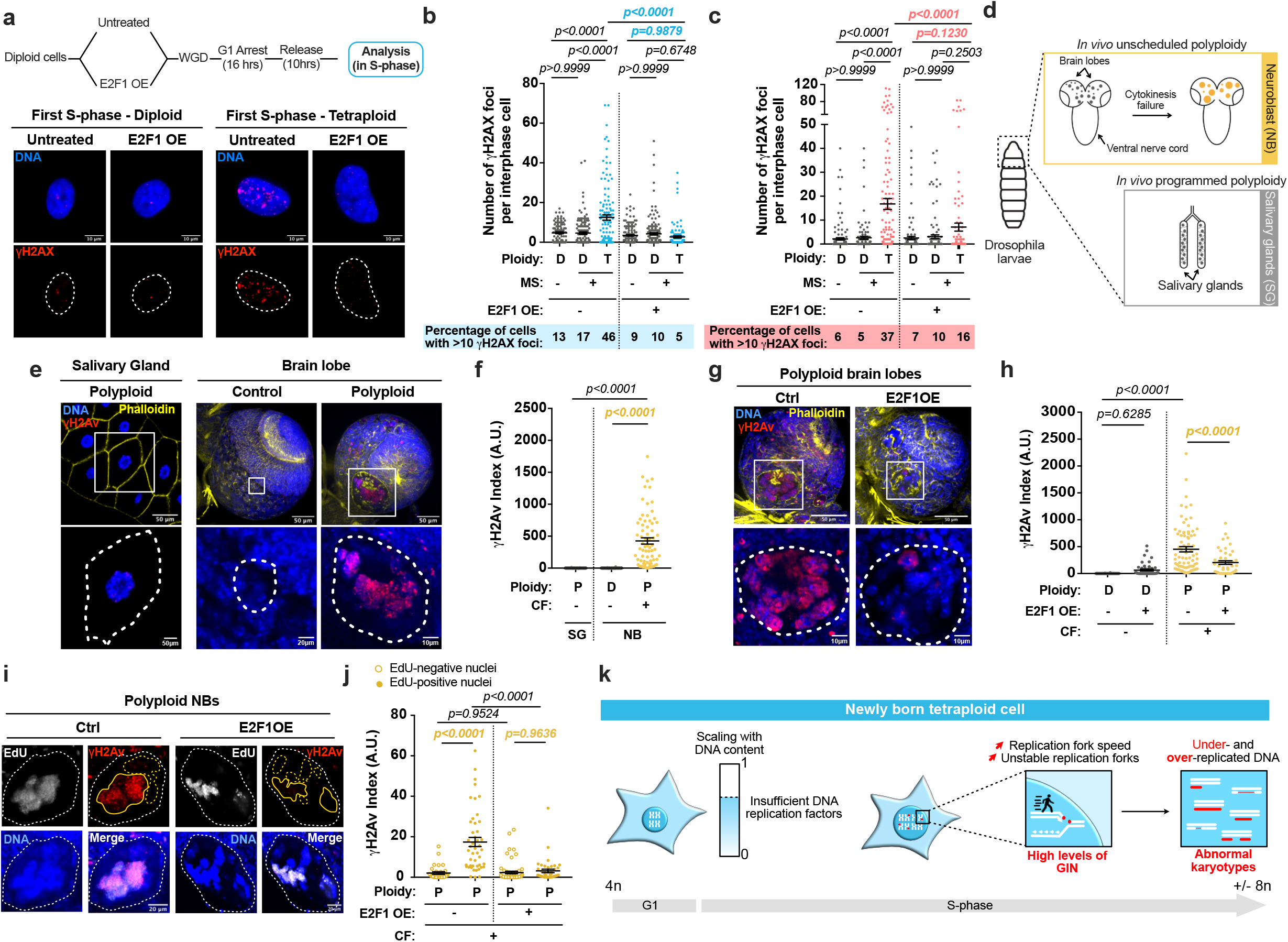
Increased E2F1 levels are sufficient to rescue GIN in both tetraploid cells and in unscheduled polyploid cells *in vivo*. **(a)** Upper - Schematic workflow showing the method used to overexpress E2F1 (E2F1OE). Lower - Images showing anti-γH2AX antibodies in red and DNA in blue. **(b-c)** Graphs showing the number of γH2AX foci per interphase cell in diploid (D, gray) and tetraploid (T) RPE-1 cells released in S-phase ± E2F1OE. Mean ± SEM, >100 interphase cells, three experiments. **(d)** Schematic representation of a *Drosophila* larva with the brain and the salivary glands (SGs). **(e)** Representative images of wild type SGs and control or *sqh* mutants brain lobes. γH2Av in red, membranes in yellow and DNA in blue. **(f)** γH2Av index in SGs or in diploid (D) and polyploid (P) neural stem cells (neuroblasts, NBs) from the *Drosophila* larvae brain. Mean ± SEM, >60 interphase cells, three experiments. **(g)** Images of brain lobes in control or *sqh* mutant ± E2F1OE. γH2Av in red, membranes in yellow and DNA in blue. **(h)** Graph showing the γH2Av index in NBs ± E2F1OE. Mean ± SEM, >30 interphase cells, three experiments. **(i)** Images of NBs in *sqh* mutant ± E2F1OE. The yellow dotted lines indicate EdU-negative nuclei, the yellow line indicates EdU461-positive nuclei. DNA in blue, γH2Av in red and EdU in gray. **(j)** Graph showing the γH2Av index in EdU-negative and -positive nuclei ± EdU. Mean ± SEM, >30 interphase cells, three experiments. **(k)** One single S-phase generates genetic instability in tetraploid cells. The white dotted lines indicate the nuclear **(a)** or cell area **(e,g** and **i)**. **(b, c, f, h** and **j)** ANOVA test (one-sided).

A key prediction of our findings is that unscheduled polyploid *Drosophila* interphase neuroblasts (NBs) ^32^ should also accumulate high levels of DNA damage *in vivo*. Indeed, the γH2Av index (methods) was higher in polyploid NBs compared to diploid NBs or to programmed polyploid salivary gland cells, which normally accumulate very large ploidies ^33^ (Fig. 4d-f). We tested the effect of E2F1OE in polyploid NBs and found that this was sufficient to decrease substantially DNA damage levels *in vivo*. Further, we noticed that DNA damage was mainly restricted to EdU^+^ nuclei (Fig. 4g-j and Extended data Fig. 10d-f). Taken together, these data show that *in vivo* unscheduled polyploidy is a source of DNA damage and GIN in replicating cells, which can be inhibited by increased E2F1 levels.

Since WGDs are quite frequent in human tumors, which have high levels of GIN ^1, 2, 34^, a prediction of our findings is that these tumors must cope with increased DNA damage levels and therefore upregulate the DDR pathway. We performed Gene set enrichment analysis using cohorts of tetraploid and diploid lung, bladder and ovarian tumors annotated in ^35^. Interestingly, this revealed an enrichment for DNA repair pathways in all tetraploid tumors when compared to diploid tumors (Extended data Fig.10h). These results suggest an increased requirement for the DDR in tumors with WGD.

## Discussion

Here, we analyzed the initial defects following WGD and identified a very early window of high GIN that could promote acquisitions of multiple mutations, making it possible to bypass cell cycle controls while promoting tetraploid cell survival. Our results are consistent with a model where tetraploid cells transit through the first cell cycle while lacking the capacity to support faithful replication of increased DNA content (Fig. 4k) (Supplementary discussion).

It is tempting to propose that in non-physiological conditions, such as those studied here, newly born tetraploids do not “feel” the increase in DNA content and therefore cannot adapt G1 duration or protein content in order to replicate a 4N genome. It will be interesting to identify the molecular mechanisms that promote ploidy increase while maintaining genetic stability and cell homeostasis to understand how tetraploid cancers, as well as tetraploids arising during evolution adapted to the new cellular state.

## Supporting information

Ext data VIdeo 1

Ext data VIdeo 2

Ext data VIdeo 3

Ext data VIdeo 4

Ext data VIdeo 5

Ext data VIdeo 6

SI Guide

## Acknowledgments

R.B dedicates this work to the memory of J.E. Rebelo, who will be deeply missed. The authors acknowledge the Cell and Tissue Imaging platform (PICT-IBiSA), member of the French National Research Infrastructure France-BioImaging (ANR10-INBS-04) and the Nikon Imaging center from Institut Curie for microscopy. We thank L. Guyonnet, A. Chipont and C. Guerrin and C. Roulin from the Cytometry and RADEXP platforms of Institut Curie. We thank V. Marthiens, S. Lambert, E. Schwob, W. Marshall, G. Almouzni, J.S. Hoffmann, D. Fachinetti, M. Budzyk, F. Edwards, G. Fantozzi, R. Salamé, A. Goupil, C. Chen, F. Perez and N. Ganem for helpful discussions and/or comments on the manuscript. This work was supported by FOR2800/STO918-7 to Z.S, ERC CoG (ChromoNumber-725907) to R.B, Institut Curie and the CNRS. The Basto lab is a member of the Cell(n)Scale Labex.

## Author contributions

S.G. and R.B. conceived the project and wrote the manuscript. DNA damage characterization and the analysis of its origins were conceived together with Z.S. S.G. did most of the experiments and data analysis presented here. M.N. did the initial observations of high levels of DNA damage in *Drosophila* polyploid NBs. R.W., A.E.T., D.C.J.S. and F.F. did the scSeq and BI analysis. K.K., S.V.B. and Z.S. contributed with DNA combing. A.S.M helped image quantifications and analysis. AS performed the 3D spheroids experiments and helped with image quantification. O.G. performed the BI analysis of ovarian TCGA tumors. N.S. and M.P. performed the quantitative phase imaging experiments and analysis and H.H. contributed with unpublished cell lines. All authors read and comment on the manuscript.

## Data availability

Methods, additional references, Nature Research reporting summaries, source data, extended data, supplementary information, peer review information; and statements of data availability are available at https://doi.org/10.6084/m9.figshare.19137323.v1.

## METHOD DETAILS

### Cell culture and generation of cell lines

#### Cell culture

Cells were maintained at 37°C in a 5% CO_2_ atmosphere. hTERT RPE-1 cells (ATCC Cat# CRL-4000, RRID:CVCL_4388) and HEK293 cells (ATCC Cat# CRL-1573, RRID:CVCL_0045) were grown in Dulbecco’s modified medium (DMEM) F12 (11320-033 from Gibco) containing 10% fetal bovine serum (GE Healthcare), 100 U/ml penicillin, 100 U/ml streptomycin (15140-122 from Gibco). BJ cells (ATCC Cat# CRL-4001, RRID:CVCL_6573) and HCT116 cells (ATCC Cat# CCL-247, RRID:CVCL_0291) were grown in Dulbecco’s modified medium + GlutaMAX (61965-026 from Gibco) containing 10% fetal bovine serum (GE Healthcare), 100 U/ml penicillin, 100 U/ml streptomycin (15140-122 from Gibco).

All cells were routinely checked for mycoplasma infection and are negative for mycoplasma infection. Identity and purity of the human cell lines used in this study were tested and confirmed using STR authentication.

#### Generation of an RPE-1 PCNA^chromo^ stable cell line

RPE-1 cells were transfected with 10µg Cell Cycle-Chromobody® plasmid (TagRFP) (From Chromotek, Planegg, Germany) using JET PRIME kit (Polyplus Transfection, 114-07) according to the manufacturer’s protocol. After 24hrs, 500µg/ml G418 (4727878001 from Sigma Aldrich) was added to the cell culture medium and then a mixed population of clones expressing PCNA chromobodies were selected.

#### Generation of an RPE-1 FUCCI or RPE-1 CCNB1^AID^ FUCCI stable cell line

To produce lentiviral particles, HEK293 cells were transfected with 4µg pBOB-EF1-FastFUCCI-Puro (86849 from Addgene, RRID:Addgene_86849) + 4µg pMD2.G (12259 from Addgene, RRID:Addgene_12259) + 4µg psPAX2 (12260 from Addgene, RRID:Addgene_12260) using a FuGENE HD Transfection Reagent (E2311 from Promega) in OptiMEM medium (51985034 from ThermoFisher). Cells were incubated at 37°C in a 5% CO2 atmosphere for 16hrs and then growth media were removed and replaced by 5 ml fresh OptiMEM. The following day, viral particles were isolated by filtering the medium containing them through a 0.45μm filter (16537 from Sartorius stedim biotech). Then, RPE-1 or RPE-1 CCNB1^AID36^ cells were incubated with viral particles in the presence of 8μg/ml polybrene (sc-134220 from Santa Cruz) at 37°C in a 5% CO_2_ atmosphere for 24hrs. RPE-1 GFP and RFP-positive cells were then collected using Sony SH800 FACS (BD FACSDiva Software Version 8.0.1). RPE-1 or RPE-1 CCNB1^AID^ clones expressing FUCCI were selected and the cell lines were established from one single clone.

pBOB-EF1-FastFUCCI-Puro was a gift from Kevin Brindle & Duncan Jodrell (Addgene plasmid # 86849; http://n2t.net/addgene:86849; RRID:Addgene_86849)^37^.

#### Generation of an RPE-1 GFP-53BP1 RFP-H2B stable cell line

This cell line was obtained as described below. Briefly, to produce lentiviral particles, HEK293 cells were transfected with 4µg pSMPUW-IRIS-Neo-H2BmRFP (Fachinetti Lab) + 4µg pMD2.G (12259 from Addgene, RRID:Addgene_12259) + 4µg psPAX2 (12260 from Addgene, RRID:Addgene_12260). Then, RPE-1 cells were incubated with viral particles and RPE-1 RFP-positive cells were collected using Sony SH800 FACS (BD FACSDiva Software Version 8.0.1). RPE-1 clones expressing RFP-H2B were selected, and the cell line was established from one single clone. Then, new lentiviral particles were produced by transfecting HEK293 cells with 4µg Apple-53BP1trunc (69531 from Addgene, RRID:Addgene_69531) + 4µg pMD2.G (12259 from Addgene, RRID:Addgene_12259) + 4µg psPAX2 (12260 from Addgene, RRID:Addgene_12260). RPE-1 RFP-H2B cells were incubated with viral particles, and RPE-1 clones expressing both RFP-H2B and GFP-53BP1 were selected using flow cytometry (FACS SH800 from Sony). The cell line was established from one single clone.

Apple-53BP1trunc was a gift from Ralph Weissleder (Addgene plasmid # 69531; http://n2t.net/addgene:69531; RRID:Addgene_69531) ^38^.

#### Generation of an RPE-1 shp53 stable cell lines

This cell line was obtained as described below. Briefly, to produce lentiviral particles, HEK293 cells were transfected with 4µg shRNA p53-puromycin (Fachinetti Lab) + 4µg pMD2.G (12259 from Addgene, RRID:Addgene_12259) + 4µg psPAX2 (12260 from Addgene, RRID:Addgene_12260). Then, RPE-1 cells were incubated with viral particles. After 24hrs, 5µg/ml puromycin (A1113803 from Gibco) was added to the cell culture medium and then a mixed population of clones expressing p53 shRNA was selected.

### Treatments

#### Induction of tetraploidy in human cell lines

To induce mitotic slippage, cells were incubated with DMSO (D8418 from Sigma Aldrich) or with 50µM monastrol (S8439 from Selleckchem) + 1µM MPI-0479605 (S7488 from Selleckchem) for at least 2hrs. Alternatively, CCNB1 depletion in RPE CCNB1^AID^ cells was induced as described before ^36^. Briefly, cells were treated with 2µg/ml doxycycline (D3447 from Sigma Aldrich) + 3µM asunaprevir (S4935 from Selleckchem) for 2hrs. Then, 500 µM auxin (I5148 from Sigma Aldrich) was added to the cell culture medium for at least 4hrs. In the figures, MS was induced by the combination of monastrol + MPI-0479605 treatment except for the following figures: Figures 2i; 3a-h, 3j-o; Extended data Fig. 2a-b; 7a and d; 8d-h; 9k, in which MS was induced by CCNB1 depletion.

To induce cytokinesis failure, cells were incubated with 10µM genistein (G6649 from Sigma Aldrich) for at least 2hrs. Alternatively, cell were incubated with 0.75µM Dihydrocytochalasin D (DCD; D1641 from Sigma-Aldrich) or with 5µM Latrunculin (L5288 from Sigma-Aldrich) for 1hr. In the figures, CF was induced by genistein treatment except for the following figures: Extended data Fig. 6j-k, in which CF was induced by DCD treatment and Extended data Fig. 2c and d, in which CF was induced by latrunculin treatment.

To induce endoreplication, cells were incubated with 10µM SP600125 (S1460 from Selleckchem) for at least 2hrs. Alternatively, CCNA2 depletion in RPE CCNA2^AID^ cells was induced as described before ^36^. Briefly, cells were treated with 2µg/ml doxycycline (D3447 from Sigma Aldrich) for 2hrs. Then, 500 µM auxin (I5148 from Sigma Aldrich) + 3µM asunaprevir (S4935 from Selleckchem) was added to the cell culture medium for at least 4hrs. In the figures, EnR was induced by SP600125 treatment except for the following figures: Figure 3q; 4c; Extended data Fig. 2e-f; 3f; 5d and j; 8c, in which EnR was induced through CCNA2 depletion.

#### Cell cycle synchronization and DNA replication inhibition

Cells were treated with 1µM palbociclib (Cdk4/6 inhibitor, S1579 from Selleckchem), or with 0.5µM abemaciclib (Cdk4/6 inhibitor, S5716 from Selleckchem) or with 1µM K03861(Cdk2 inhibitor, S8100 from Selleckchem) for 16hrs to synchronize cells at G1/S transition, and were collected (indicated by “G1 arrest” in the figures). Alternatively, cells were then washed five times using PBS 1X and released in S-phase for 10hrs before being collected. To extend G1 duration cells were treated with 160nM palbociclib or with 50nM abemaciclib or with 400nM K03861 for 16hrs and were collected (indicated by “G1 lengthening” in the figures). Alternatively, cells were then washed five times using PBS 1X and released in S-phase for 10hrs before being collected.

To inhibit DNA replication, cells were released in S-phase in the presence of low doses of Aphidicolin (APH, A0781 from Sigma-Aldrich), a DNA replication polymerase inhibitor, or of PHA767491 (PZ0178 from Sigma-Aldrich), a Cdc7 inhibitor (indicated by “Release in S-phase + APH” or “Release in S-phase + PHA”, respectively, in the figures). Doses were chosen to significantly decrease EdU incorporation without affecting the levels of DNA damage.

#### Nucleoside supplementation

Cells were synchronized in G1 using 1µM palbociclib and then released in S phase (see *Cell cycle synchronization and DNA replication inhibition* section) in the presence of nucleosides at the following concentrations: dC 7,3mg/L (D0776 from Sigma Aldrich); dG 8,5mg/L (D0901 from Sigma Aldrich); dU 7,3mg/L (D5412 from Sigma Aldrich); dA 8mg/L (D8668 from Sigma Aldrich) and dT 2,4mg/L (T1895 from Sigma Aldrich) (+ in the figures) or dC 14,6mg/L; dG 17mg/L; dU 14,6mg/L; dA 16mg/L and dT 4,8mg/L (++ in the figures).

#### Treatments

The drugs were used at the following concentrations: Auxin (I5148 from Sigma), 500µM; Doxycycline (D3447 from Sigma), 2µg/ml; Asunaprevir (S4935 from Selleckchem), 3µM; Monastrol (S8439 from Selleckchem), 50µM; MPI-0479605 (S7488 from Selleckchem), 1µM; Genistein (G6649 from Sigma), 10µM; SP600125 (S1460 from Selleckchem), 10µM; Abemaciclib (S5716 from Selleckchem), 50nM or 0,5µM; K03861 (S8100 from Selleckchem), 400nM or 1µM; Palbociclib (S1579 from Selleckchem), 120nM or 1µM; Aphidicolin (A0781 from Sigma), 0,4µM or 1µM; Hydroxyurea (S1896 from Selleckchem), 2mM; PHA767491 (PZ0178 from Sigma), 1µM; RO3306 (217699 from Calbiochem), 10µM; Dihydrocytochalasin D (D1641 from Sigma), 0,75µM; Latrunculin B (L5288 from Sigma), 5µM; 5’-Chloro-2’-deoxyuridine (CIdU) (C6891 from Sigma), 100µM; 5’-Iodo-2’-deoxyuridine (IdU) (I7125 from Sigma), 100µM.

### Fly husbandry and fly stocks

Flies were raised on cornmeal medium (0.75% agar, 3.5% organic wheat flour, 5.0% yeast, 5.5% sugar, 2.5% nipagin, 1.0% penicillin-streptomycin and 0.4% propionic acid). Fly stocks were maintained at 18°C. Crosses were carried out in plastic vials and maintained at 25°C. Stocks were maintained using balancer inverted chromosomes to prevent recombination. Stocks used in this study: *sqh*^1 39^, *pavarotti* RNAi (Pav ^RNAi^) (BL#42573 from Bloomington *Drosophila* Stock Center, Indiana University, IN, USA) ^32^, UAS-E2F1 (F001065 from FlyORF, Zurich, Switzerland) and UAS-Rb (BL# 50746 from Bloomington Drosophila Stock Center, Indiana University, IN, USA).

In all experiments, larvae were staged to obtain comparable stages of development. Egg collection was performed at 25°C for 24hrs. After development at 25°C, third instar larvae were used for dissection.

### Immunofluorescence microscopy and antibodies

#### Preparation and imaging of human cells

Cells were plated on cover slips in 12-well plates and treated with the indicated drugs. To label cells, they were fixed using 4% of paraformaldehyde (15710 from Electron Microscopy Sciences) + Triton X-100 (2000-C from Euromedex) 0,1% in PBS (20 min at 4°C). Then, cells were washed three times using PBS-T (PBS 1X + 0,1% Triton X-100 + 0,02% Sodium Azide) and incubated with PBS-T + BSA (04-100-812-C from Euromedex) 1% for 30 min at RT. After three washes with PBS-T + BSA, primary and secondary antibodies were incubated in PBS-T + BSA 1% for 1hr and 30 min at RT, respectively. After two washes with PBS, cells were incubated with 3 μg/ml DAPI (4’,6-diamidino-2-phenylindole; D8417 from Sigma Aldrich) for 15 min at RT. After two washes with PBS, slides were mounted using 1.25% n-propyl gallate (Sigma, P3130), 75% glycerol (bidistilled, 99.5%, VWR, 24388-295), 23.75% H2O. Images were acquired on an upright widefield microscope (DM6B, Leica Systems, Germany) equipped with a motorized XY and a 40X objective (HCX PL APO 40X/1,40-0,70 Oil from Leica). Acquisitions were performed using Metamorph 7.10.1 software (Molecular Devices, USA) and a sCMOS camera (Flash 4V2, Hamamatsu, Japan). Stacks of conventional fluorescence images were collected automatically at a Z-distance of 0.5 µm (Metamorph 7.10.1 software; Molecular Devices, RRID:SCR_002368). Images are presented as maximum intensity projections generated with ImageJ software (RRID:SCR_002285).

#### Whole mount tissue preparation and imaging of Drosophila larval brains

Brains or Salivary glands from third instar larvae were dissected in PBS and fixed for 30 minutes in 4% paraformaldehyde in PBS. They were washed three times in PBST 0.3% (PBS, 0.3% Triton X-100 (T9284, Sigma), 10 minutes for each wash) and incubated for several hrs in agitation at room temperature (RT) and O/N at 4°C with primary antibodies at the appropriate dilution in PBST 0.3%. Tissues were washed three times in PBST 0.3% (10 minutes for each wash) and incubated O/N at 4°C with secondary antibodies diluted in PBST 0.3%. Brains and salivary glands were then washed two times in PBST 0.3% (30 minutes for each wash), rinsed in PBS and incubated with 3 μg/ml DAPI (4’,6-diamidino-2-phenylindole; D8417 from Sigma Aldrich) at RT for 30min. Brains and salivary glands were then washed in PBST 0.3% at RT for 30 minutes and mounted on mounting media. A standard mounting medium was prepared with 1.25% n-propyl gallate (Sigma, P3130), 75% glycerol (bidistilled, 99.5%, VWR, 24388-295), 23.75% H2O.

Images were acquired on a spinning disk microscope (Gataca Systems, France). Based on a CSU-W1 (Yokogawa, Japan), the spinning head was mounted on an inverted Eclipse Ti2 microscope equipped with a motorized XY Stage (Nikon, Japan). Images were acquired through a 40X NA 1.3 oil objective with a sCMOS camera (Prime95B, Photometrics, USA). Optical sectioning was achieved using a piezo stage (Nano-z series, Mad City Lab, USA). The Gataca Systems’ laser bench was equipped with 405, 491 and 561 nm laser diodes, delivering 150 mW each, coupled to the spinning disk head through a single mode fibre. Multi-dimensional acquisitions were performed using Metamorph 7.10.1 software (Molecular Devices, USA). Stacks of conventional fluorescence images were collected automatically at a Z-distance of 1.5 µm (Metamorph 7.10.1 software; Molecular Devices, RRID:SCR_002368). Images are presented as maximum intensity projections generated with ImageJ software (RRID:SCR_002285).

#### Primary and secondary antibodies

Primary and secondary antibodies were used at the following concentrations: Guinea pig anti CEP192 antibody (1/500; Basto lab)^40^, rabbit anti-beta catenin (1/250; C2206 from Sigma-Aldrich, RRID:AB_476831), mouse anti-gamma H2A.X phospho S139 (1/1000; ab22551 from Abcam, RRID:AB_447150), mouse anti-XRCC1 (1/500; ab1838 from Abcam, RRID:AB_302636), rabbit anti-Rad51 (1/500; ab133534 from Abcam, RRID:AB_2722613), mouse anti-KU80 (1/200; MA5-12933 from ThermoFisher, RRID:AB_10983840), rabbit anti-FANCD2 (1/150; NB100-182SS from Novusbio, RRID:AB_1108397), mouse anti-53BP1 (1/250; MAB3802 from Millipore, RRID:AB_2206767), rabbit anti-γH2Av (1/500; 600-401-914 from Rockland; RRID: AB_11183655), Alexa Fluor® 647 Phalloidin (1/250; A22287 from ThermoFisher Scientific, RRID:AB_2620155), goat anti-Rabbit IgG (H+L) Highly Cross-Adsorbed Secondary Antibody, Alexa Fluor 647 (1/250; A21245 from ThermoFisher, RRID:AB_2535813), Goat anti-Guinea Pig IgG (H+L) Highly Cross-Adsorbed Secondary Antibody, Alexa Fluor 488 (1/250; A11073 from ThermoFisher, RRID:AB_253411), Goat anti-Mouse IgG (H+L) Cross-Adsorbed Secondary Antibody, Alexa Fluor 546 (1/250, A11003 from ThermoFisher, RRID:AB_2534071), Goat anti-Rabbit IgG (H+L) Highly Cross-Adsorbed Secondary Antibody, Alexa Fluor 546 (1/250; A-11035 from Thermo Fisher Scientific, RRID:AB_2534093).

### Quantitative analysis of DNA damage

#### Analysis of Drosophila NBs and of 3D spheroids

Quantitative analysis of DNA damage was carried out as previously described ^32^. In brief, DNA damage was assessed using a γH2Av primary antibody for *Drosophila* and with γH2AX for 3D spheroids detected with an Alexa Fluor secondary antibody. Confocal volumes were obtained with optical sections at 1.5 µm intervals. Image analysis was performed using Fiji and a custom plugin developed by QUANTACELL. After manual segmentation of the nuclei, a thresholding operation was used to determine the percentage of γH2Av/γH2AX positive pixels (coverage) and their average intensity in a single projection. Coverage and intensity were multiplied to obtain the γH2Av/γH2AX index. The threshold used to detect and quantify the γH2Av index in polyploid NBs does not detect any damage in SGs. However, it is important to mention that in a fraction of these cells, γH2Av dots (small and of low FI) can be occasionally seen.

#### Analysis of 2D human cell lines

For DNA damage quantification, the signals obtained in cultured cells were different from the signals found in *Drosophila* NBs. To asses DNA damage in human cells, we used an ImageJ software-based plugin developed by QUANTACELL, where γH2AX signals were measured using z-projection stacks after thresholding. Nuclear size, DAPI intensity, the number of γH2AX foci, γH2AX FI and the percentage of nuclear coverage by γH2AX signal were obtained for each nucleus.

### Time lapse microscopy

Cells were plated on a dish (627870 from Dutscher) and treated with the indicated drugs. Images were acquired on a spinning disk microscope (Gataca Systems, France). Based on a CSU-W1 (Yokogawa, Japan), the spinning head was mounted on an inverted Eclipse Ti2 microscope equipped with a motorized XY Stage (Nikon, Japan). Images were acquired through a 40X NA 1.3 oil objective with a sCMOS camera (Prime95B, Photometrics, USA). Optical sectioning was achieved using a piezo stage (Nano-z series, Mad City Lab, USA). Gataca Systems’ laser bench was equipped with 405, 491 and 561 nm laser diodes, delivering 150 mW each, coupled to the spinning disk head through a single mode fiber. Laser power was chosen to obtain the best ratio of signal/background while avoiding phototoxicity. Multi-dimensional acquisitions were performed using Metamorph 7.10.1 software (Molecular Devices, USA). Stacks of conventional fluorescence images were collected automatically at a Z-distance of 0.5 µm (Metamorph 7.10.1 software; Molecular Devices, RRID:SCR_002368). Images are presented as maximum intensity projections generated with ImageJ software (RRID:SCR_002285), from stacks deconvolved with an extension of Metamorph 7.10.1 software.

### 3D cultures

#### Mitotic slippage on 3D cultures

To generate spheroids, 500 cells per well were seeded into 96 ultra low attachment well plates (7007 from Corning) in presence of DMSO (D8418 from Sigma Aldrich) or with 50µM monastrol (S8439 from Selleckchem) and 1µM MPI-0479605 (S7488 from Selleckchem). Plates were spin down at 200g for 3min, to allow spheroid formation, and incubated for 24hrs at 37°C.

#### Immunostaining

Spheroids were collected and washed quickly with PBS prior fixation using 4% of paraformaldehyde (15710 from Electron Microscopy Sciences) in PBS for 40min. Then, spheroids were permeabilized for 5 min using Triton X-100 (2000-C from Euromedex) 0,3% in PBS and blocked for 30min using Blocking Buffer (BB= PBS 1X + 0,3% Triton X-100 + 0,02% Sodium Azide + 3% BSA). Aggregates were incubated with primary antibodies diluted into BB O/N. After 3 washes using BB, spheroids were incubated with secondary antibodies in BB for 3hrs. Cells were then washed several times for 2hrs in BB and mounted on glass with EverBrite (Biotium). For primary and secondary antibodies see *Immunofluorescence microscopy and antibodies* section.

#### Imaging and DNA damage analysis

Spheroids were imaged using an inverted scanning laser confocal (Nikon A1RHD25) equipped with a 100x CFI Plan Apo Lambda S Sil objective (NA 1.35). Z-stacks were acquired every 0.3 um. Diploid and tetraploid cells were distinguished using cell and nuclear size and centrosome number. Then, quantitative analysis of DNA damage was carried out (see Q*uantitative analysis of DNA damage* section).

### EdU staining

EdU incorporation into DNA was visualized with the Click-it EdU imaging kit (C10338 from Life Technologies), according to the manufacturer’s instructions. For human cell lines, EdU was used at a concentration of 1µM (Extended data Fig. 6e and Extended data Fig. 9h) or 10 µM (Extended data Fig. 5g and h) for the indicated time. Cells were incubated with the Click-it reaction cocktail for 15 minutes. EdU incorporation in polyploid NBs was done as previously described ^32^ with a pulse of 2hrs before fixation.

### Comet assay

Comet assays were performed using Single Cell Gel Electrophoresis Assay kit (4250-050-ES from Trevigen) according to the manufacturer’s instructions. Comets were then imaged using an inverted Eclipse Ti-E Nikon videomicroscope equipped with a 40x CFI Plan Fluor objective. Images were analyzed with OpenComet plugin on Fiji. Based on the comet DNA content of DMSO treated cells, a manual threshold was applied to identify diploid from tetraploid cells. The same threshold was applied on the cells treated for mitotic slippage.

### FACS sorting of diploid and tetraploid cells

A mix of diploid and tetraploid cells (see *Induction of tetraploidy in human cell lines* section) were incubated with 2µg/ml Hoescht 33342 (94403 from Sigma Aldrich) for 1hr at 37°C, 5% CO_2_. Then, a single cell suspension was generated. Cells were washed using PBS 1X, the supernatant was removed and cells were resuspended in a cold cell culture medium at 1×10^7^ cell per ml and kept at 4°C during all the experiments. FACS sorting was performed using Sony SH800 FACS (BD FACSDiva Software Version 8.0.1). Compensation was performed using the appropriate negative control samples. Experimental samples were then recorded and sorted using gating tools to select the populations of interest. RFP+ / GFP-negative cells (G1 cells) were first selected. Then, in this population, DNA content was used to segregate diploid (2n) and tetraploid (4n) G1 cells (*see* Extended data Fig 8d). Once gates have been determined, the same number of diploid and tetraploid G1 cells were sorted into external collection tubes. The number of cells was then checked using a cell counter and the same number of diploid an tetraploid cells were collected for western blot analysis (see *Western Blot analysis, subcellular fractionation and antibodies* section). In parallel, post-sort analysis was performed to determine the purity of the sorted populations (*see* Extended data Fig. 8e).

### Cell cycle analysis and measure of RNA levels by flow cytometry

Cells were detached by treatment with Accutase (Sigma), immediately washed in 1x PBS, fixed in 2ml 70% ethanol and stored at −20°C overnight. They were then washed in 1x PBS and staining buffer (554656 from BD Pharmingen).

For cell cycle analysis, DNA content was visualized by incubating the cells with 2µg/ml Hoescht 33342 (94403 from Sigma Aldrich) in staining buffer for 15 min at room temperature. Alternatively, to measure RNA levels, cells were incubated with 2µg/ml Hoescht 33342 + Pyronin 4µg/ml (sc-203755A from Santa Cruz) in a staining buffer for 20 min at room temperature. Flow cytometry analysis was done using LSRII (from BD Biosciences), by analyzing 10 000 cells per condition. Data were then analyzed with FlowJo 10.6.0 software (Tree Star Inc.).

### E2F1 overexpression (OE)

RPE-1 cells were transfected using 0.25µg pCMVHA E2F1 (24225 from Addgene, RRID: Addgene_24225) with a JET PRIME kit (Polyplus Transfection, 114-07) according to the manufacturer’s protocol. 5hrs later, cells were incubated with DMSO (D8418 from Sigma Aldrich) or with 50µM monastrol (S8439 from Selleckchem) + 1µM MPI-0479605 (S7488 from Selleckchem) to generate tetraploid cells. After 2hrs, DMSO or 1µM palbociclib (S1579 from Sellechem) were added to the cell culture medium for 16hrs. Cells were then fixed in G1 (T0) or washed five times using PBS and released in S-phase and fixed after 10hrs (T10). The immunofluorescence protocol is described in the corresponding section.

pCMVHA E2F1 was a gift from Kristian Helin (Addgene plasmid # 24225; http://n2t.net/addgene:24225; RRID:Addgene_24225) ^41^.

### Western Blot analysis, subcellular fractionation and antibodies

#### Western Blot

For a whole-cell extract, cells were lysed in 8 M urea, 50 mM Tris HCl, pH 7.5 and 150 mM β-mercaptoethanol (161-0710 from Bio-Rad), sonicated and heated at 95°C for 10 minutes. For chromatin bound fractions, cells were prepared using the Subcellular Protein Fractionation Kit for Cultured Cells (78840 from ThermoFisher Scientific), according to the manufacturer’s instructions. Then, samples (equivalent of 2 x 10^5^ cells) were subjected to electrophoresis in NuPAGE Novex 4–12% Bis-Tris pre-cast gels (NP0321 from Life Technologies). The same number of cells (see *FACS sorting of diploid and tetraploid cells* section) were loaded for diploid and tetraploid conditions, allowing us to compare one diploid cell with one tetraploid cell. Protein fractions from the gel were electrophoretically transferred to PVDF membranes (PVDF transfer membrane; RPN303F from GE). After 1hr saturation in PBS containing 5% dry non-fat milk and 0.5% Tween 20, the membranes were incubated for 1hr with a primary antibody (*see below*) diluted in PBS containing 5% dry non-fat milk and 0.5% Tween 20. After three 10-min washes with PBS containing 0.5% Tween 20, the membranes were incubated for 45 min with a 1/2 500 dilution of peroxidase-conjugated antibody (*see below*). Membranes were then washed three times with PBS containing 0.5% Tween 20, and the reaction was developed according to the manufacturer’s specifications using ECL reagent (SuperSignal West Pico Chemiluminescent Substrate; 34080 from Thermo Scientific).

The background-adjusted volume intensity was calculated and normalized using a H2B signal (H2B was used as a readout of DNA content) for each protein, using Image Lab software version 6.0.1, Bio-Rad Laboratories. All the original uncropped blots (gel source data) are presented in Supplementary Figure 1.

#### Primary and secondary antibodies were used at the following concentrations

Mouse anti-αtubulin (1/5000; T9026 from Sigma, RRID:AB_477593), mouse anti-CDC45 (1/100; sc-55569 from Santa Cruz Biotechnology, RRID:AB_831146), rabbit anti-PCNA (1/500; sc56 from Santa Cruz, RRID:AB_628110), rabbit anti-Actin (1/2000; A5060 from Sigma-Aldrich, RRID:AB_476738), mouse anti-H2B (1/1000; sc-515808 from Santa Cruz Biotechnology), mouse anti-ORC1 (1/100; sc-398734 from Santa Cruz Biotechnology), mouse anti-MCM2 (1/500; 610701 from BD Biosciences, RRID:AB_398024), mouse anti-E2F1 (1/2000; sc251 from Santa Cruz, RRID:AB_627476), mouse anti-CDC6 (1/500; sc-9964 from Santa Cruz, RRID:AB_627236), rabbit anti-CDT1 (1/500; 8064S from Cell Signaling, RRID:AB_10896851), rabbit anti-Treslin (1/500; A303-472A from Betyl, RRID:AB_10953949), Goat anti-Rabbit IgG (H+L) Cross-Adsorbed Secondary Antibody, HRP (1/2500; G21234 from ThermoFisher, RRID:AB_2536530), Peroxidase AffiniPure Goat Anti-Mouse IgG (H+L) (1/2500; 115-035-003 from Jackson ImmunoResearch, RRID:AB_10015289).

### 3D reconstruction and analysis on Imaris

3D Videos (see *time lapse microscopy* section) were imported into *Imaris* software v.9.6.0 (Bitplane, RRID:SCR_007370). For chosen cells, the module “Spot tracking” of *Imaris* v.9.6.0 was used to detect the foci, as spots of diameter 0.5 µm in the XY-direction and 1µm in Z-direction (modelling PSF elongation). Because the volume of the foci changes in time, the option “Enable growing regions” was used. In each video, the threshold was chosen on the brightest frame (to detect a maximum of the correct spots) and then applied to the whole video. For each cell, at each time point, the number of spots and volumes were recorded. To determine DNA replication timing, we quantified the signal of PCNA fluorescence intensity (FI) in the nucleus. This replication timing was characterized independently of any particular behavior of PCNA. As soon as PCNA FI was detected in the nucleus, t=0 (beginning of S-phase) was defined, and when PCNA FI was not detected anymore the last time point was defined (end of S-phase). For each condition, at least 10 cells (see Supplementary data 1) were studied and the statistics from *Imaris* v.9.6.0 were averaged at each time point using a MATLAB script.

### Molecular combing and antibodies

#### Molecular combing

Tetraploid HCT116 were generated by cytokinesis inhibition using 0.75 µM dihydrocytochalasin D (DCD, inhibitor of actin polymerization, D1641 from Sigma-Aldrich) for 18hrs overnight. Afterwards, the cells were washed three times with PBS and cultured in DMEM supplemented with 10% FBS and 1% Pen/Strep for additional 20hrs. Tetraploid RPE-1 and BJ cells were generated by mitotic slippage or endoreplication (see *Induction of tetraploidy in human cell lines*). Then, the cells were washed three times with PBS and cultured in DMEM supplemented with 10% FBS and 1% Pen/Strep for additional 20hrs. For each method, we determined that the proportion of tetraploid cells in the treated population is about 40-60%. Due to the presence of diploid cells in the treated population, the consequences of tetraploidization on replication fork speed, fork asymmetry and inter-origin distance (IOD) are most likely underestimated.

Diploid controls and the tetraploid-enriched population were then pulse-labelled with 0.1 mM CIdU and 0.1 mM IdU for 30min and 100 000-300 000 cells per condition were collected for further analysis. The DNA was extracted from cells and prepped following the manufacturer’s instructions using the FiberPrep® DNA Extraction Kit (Genomic Vision, Bagneux, France). Subsequently, the prepped DNA was stretched onto coated glass coverslips (CombiCoverslips™, Genomic Vision, Bagneux, France) by using the FiberComb Molecular Combing System (Genomic Vision, Bagneux, France). The Labeling was performed with antibodies against ssDNA, IdU and CldU using the Replication Combing Assay (RCA) (Genomic Vision, Bagneux, France). The imaging of the prepared cover slips was carried out by Genomic Vision (Bagneux, France) and analysed using the FiberStudio® 2.0.1 Analysis Software by Genomic Vision. Replication speed was determined by measuring the combined length of the CldU and IdU tracks. Fork asymmetry was determined by measuring symmetry of the CldU and IdU incorporation by the forks (the length of the first track -CldU - is compared to the length of the second track - IdU). IOD was determined by measuring distance between two origins on the same fibers.

#### Antibodies were used at the following concentrations

Rabbit anti-ssDNA (1/5; 18731 from IBL International, RRID: AB_494649), Rat anti-CldU (1/10; Ab6326 from Abcam, RRID:AB_2313786), Mouse anti-IdU (1/10; 555627 from BD Biosciences, RRID:AB_10015222), mouse Alexa Fluor 647 Donkey (1/25; JIM-715-605-151 from Biozol), Rat Alexa Fluor 594 Donkey (1/25; JIM-712-585-153 from Biozol), Rabbit Brilliant Violet 480 Donkey (1/25; 711-685-152 from Jackson Immuno Research, RRID: AB_2651109).

### Quantitative phase imaging and measurements

Cells were plated on glass-bottom dishes coated with 50 µg/ml Fibronectin for 1hr and rinsed, and trypsinised cells were plated at a concentration of 1.5×10^6^ cells/ml. The cells used for the experiments were seeded in T-25 dishes at a concentration of 0.7×10^6^ cells/ml 2 days before the actual experiment. On the day of the experiment, the cells were detached with EDTA (versene), and plated at a concentration of 1.5×10^6^ cells/ml. For inducing tetraploidy, cells were treated with 2µg/ml doxycycline (D3447 from Sigma Aldrich) for 2hrs. Then, 500 µM auxin (I5148 from Sigma Aldrich) + 3µM asunaprevir (S4935 from Selleckchem) was added to the cell culture medium for at least 4hrs. The cells were then imaged for 35hrs every 20 minutes to track them throughout their cell cycle.

The cell cycle state was indicated by the FUCCI system; G1 cells express Cdt1-RFP while S/G2 cells express hGeminin-GFP and mitosis was indicated by the nuclear envelope break down (NEBD) with geminin being present through the cells ^42^. To quantify the fluorescence of geminin in the nucleus, first a background subtraction was performed on the images. An ROI was used to define an area containing the background fluorescence in the image. An average value of the ROI was then subtracted from all the frames. Subsequently, a ROI was drawn as close as possible to the cell, and then the mean gray value was measured across all the frames. This helped identify the frames of birth and G1/S transition during the cell cycle.

A detailed protocol for the mass measurement with phasics camera is available in ^43, 44^. Images were acquired by a Phasics camera every 20min for 35hrs for the duration of the experiment. To obtain the reference image, 32 empty fields were acquired on the dish and a median image was calculated. This reference image was subtracted from the interferograms (images acquired by phasics) by custom written MATLAB scripts to measure the optical path difference. They were then processed to calculate the phase, intensity and phase cleaned images (the background set to 1000 and the field cropped to remove edges). Background normalization was performed using a gridfit method, and a watershed algorithm was used to separate cells which came in contact with each other. Mass was calculated by integrating the intensity of the whole cell.

### Sequencing and AneuFinder analysis

A mixed population of diploid and tetraploid RPE-1 CCNB1^AID^ FUCCI cells were synchronized in G1 using 1µM palbociclib (S1579 from Selleckchem) for 16hrs or released in S-phase for 20hrs in the presence of 10µM RO3306 (217699 from Calbiochem) in order to block cells in the subsequent G2/M. G1 and G2/M diploid and tetraploid cells were then isolated using cell sorting (see *FACS sorting of diploid and tetraploid cells* section) and collected in a 96-well plate.

Sequencing was performed using a NextSeq 500 machine (Illumina; up to 77 cycles; single end). The generated data were subsequently demultiplexed using sample-specific barcodes and changed into fastq files using bcl2fastq (Illumina; version 1.8.4). Reads were afterwards aligned to the human reference genome (GRCh38/hg38) using Bowtie2 (version 2.2.4;^45^. Duplicate reads were marked with BamUtil (version 1.0.3; ^46^. The aligned read data (bam files) were analyzed with a copy number calling algorithm called AneuFinder (https://github.com/ataudt/aneufinder); ^47^. Following GC correction and blacklisting of artefact-prone regions (extreme low or high coverage in control samples), libraries were analyzed using the dnacopy and edivisive copy number calling algorithms with variable width bins (average binsize = 1 Mb; step size = 500 kb). The G1 samples were analyzed with an euploid reference (van den Bos et al., 2016). The G1 samples were used as a reference for the analysis of the G2/M samples (G1 diploid for G2/M diploid and G1 polyploid for G2/M polyploid). Aneuploid libraries were not used as a reference and blacklists were constructed using the example from Bioconductor as a guideline. The RPE-1 diploid G1 sample (2n) was analyzed with the standard version of AneuFinder (from Bioconductor) while the other samples were analyzed with the developer version of AneuFinder (from GitHub; 4n and 8n samples). The ground ploidy for these samples was constrained between 3.5 and 4.5 (4n samples) or between 7.5 and 8.5 (8n samples; parameters: min.ground.ploidy and max.ground.ploidy). Results were afterwards curated by requiring a minimum concordance of 95 % (2n sample) or 90% (4n and 8n samples) between the results of the two algorithms. Libraries with on average less than 10 reads per chromosome copy of each bin (2-somy: 20 reads, 3-somy: 30 reads, etc.) were discarded. This minimum number of reads comes down to roughly 60,000 for a diploid genome in G1 phase (2n) up to 240,000 for a polyploid genome in G2/M phase (8n). Analysis of the BJ samples showed aberrations (wavy patterns) that resulted in wrongly called segments with a copy number which is either one higher or one lower than the expected state (when euploid). The means of the read counts (read counts of the bins) of these states were too close to the mean of the expected state (e.g. mean 5-somy too close to mean 4-somy; 4n sample; see Supplementary Methods 1). When more than 1 % of the genome was classified as such (e.g. more than 1 % 5-somy), a non-rounded version of the copy number of the state was calculated using the mean of the expected state (ploidy of euploid sample) as a reference:

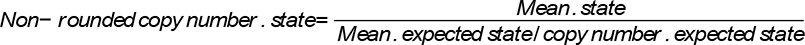

Example 5-somy (4n sample):

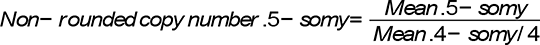

This was done in order to quantify the distance between the two states. The values are typically between −0.5 and +0.5 of the state under consideration (e.g. 5-somy; between 4.5 and 5.5), which will result in a rounded value equal to the state. The libraries with aberrations have typically a deviation of 0.25 and more from the expected value (see Supplementary Methods 1). Libraries that showed a deviation of more than 0.25 were therefore discarded (For 5-somy; a value lower than 4.75 or higher than 5.25). By applying this cutoff, we eliminated libraries that clearly showed this aberration (see Supplementary Methods 1) while preserving true aneuploid libraries (see Supplementary Methods 1). This specific method was only used for the BJ samples.

### Gene Set Enrichment Analysis with TCGA PanCancer data

Gene Set Enrichment Analysis (GSEA) was performed using GSEA software v.4.2.1 ^48, 49^. The normalized mRNA Expression (Illumina HiSeq_RNASeqV2, RSEM) from pan cancer studies were downloaded from https://www.cbioportal.org/: detailed information about RNA sequencing experiment and tools used can be found at the NCI’s Genomic Data Commons (GDC) portal https://gdc.cancer.gov. The ploidy status for bladder urothelial carcinoma (156 near-diploid and 200 near-tetraploid samples), Lung adenocarcinoma (205 near-diploid and 240 near-tetraploid samples), and ovarian serous cystadenocarcinoma (116 near-diploid and 130 near-tetraploid samples) were extracted from ^35^. In addition to ranked list of genes and ploidy status, we use gene sets derived from the GO Biological Process ontology to assess significant pathway enrichment between near -diploid and near tetraploid tumors in GSEA tool. Gene Set Enrichment Analysis (GSEA) is a computational method that determines whether an *a priori* defined set of genes shows statistically significant, concordant differences between two biological states (e.g. phenotypes). ^48^ fully describes the algorithm based on the calculation of an Enrichment score (ES), the estimation of significance level of ES (nominal P Value) and adjustment for multiple hypothesis testing (ES normalization, FDR calculation…).

### Quantifications and statistical analysis

#### Quantifications

Image analysis and quantifications were performed using Image J software V2.1.0/1.53c, https://imagej.net/software/fiji/downloads. To quantify the colocalizations between two signals (Extended data Fig. 3i and m, Extended data Fig. 4g and j) we calculated the Manders coefficient using *JACOP* plugin with Image J V2.1.0/1.53c software. We determined that the colocalizations between γH2AX signal and EdU, FANCD2 or RAD51 signals are not random using an home-made based Costes randomization on nuclear area with Image J software. 1000 randomizations of the pixel positions were performed for each condition (Supplementary data 2). 3D Videos (Extended data Fig. 3c, Extended data Fig. 6c and Extended data Fig. 9c and d) were corrected using *3D correct drift* plugin with Image J V2.1.0/1.53c software to keep the cell of interest at the center of the region of interest. The nuclear area and DAPI intensity were measured using the *wand* tool with Image J V2.1.0/1.53c software. For the figures, images were processed on Image J V2.1.0/1.53c software, and mounted using Affinity Designer, https://affinity.serif.com/fr/designer/.

#### Statistical and reproducibility

At least two (n) independent experiments were carried out to generate each dataset, and the statistical significance of differences was calculated using GraphPad Prism (RRID:SCR_002798) version 7.00 for Mac, GraphPad Software, La Jolla California USA, www.graphpad.com. The statistical test used for each experiment is indicated in the figure legends. Each representative image (Fig. 2a, c; 3a, e, g, k, n; 4a; and Extended Data Fig. 2a, c, e, l; 3a, g; 4c; 5g; 6c; 9c, d; 10a) originates from a dataset composed of at least two (n) independent experiments.

## FIGURE LEGENDS FOR EXTENDED DATA FIGURES

**Extended data Figure 1:**
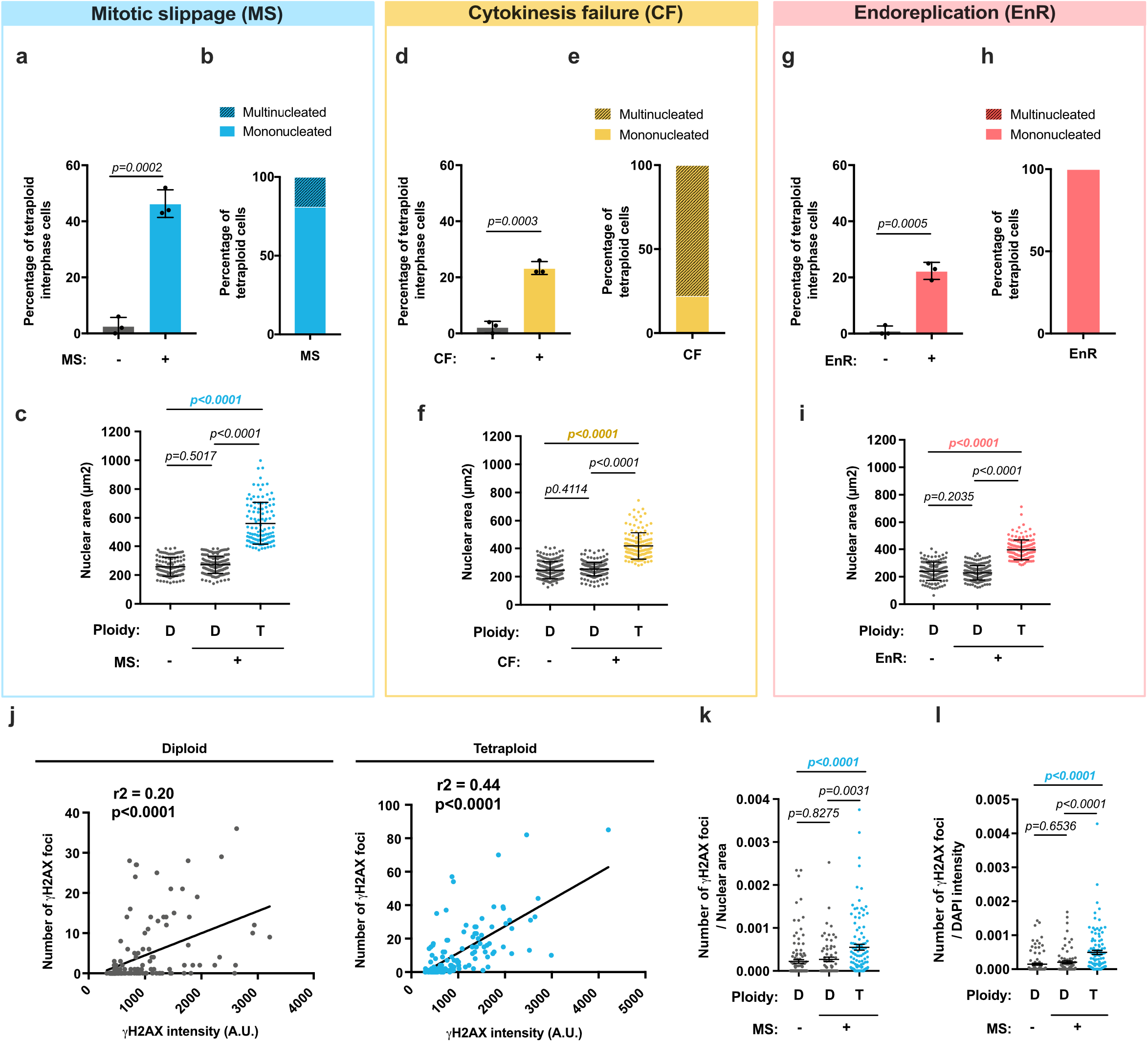
Characterization of RPE-1 cells upon WGD. **(a, d** and **g)** Graphs showing the percentage of tetraploid interphase RPE-1 cells in the indicated experimental conditions. Mean ± SEM, > 100 interphase cells were analyzed from three independent experiments. **(b, e** and **h)** Graphs showing the percentage of mono- and multinucleated RPE-1 tetraploid cells in the indicated experimental conditions. Mean ± SEM, > 100 interphase cells were analyzed from three independent experiments. **(c, f** and **i)** Graphs representing the nuclear area in diploid (D) and tetraploid (T) RPE-1 cells. Mean ± SEM, >100 interphase cells were analyzed from three independent experiments. **(j)** Graph showing the correlation between the number of γH2AX foci and γH2AX foci intensity in diploid (left panel, gray) and tetraploid (right panel, blue) RPE-1 cells induced through MS. >100 interphase cells were analyzed from three independent experiments**. (k-l)** Graphs showing the number of γH2AX foci relative to nuclear area **(k)** or DAPI fluorescence intensity (FI) **(l)** in diploid (gray) and tetraploid (blue) RPE-1 cells induced through MS. t-test (two-sided) **(a, d** and **g)**. ANOVA test (one-sided) **(c, f, i, k** and **l)**. Pearson test (two-sided) **(j)**.

**Extended data Figure 2:**
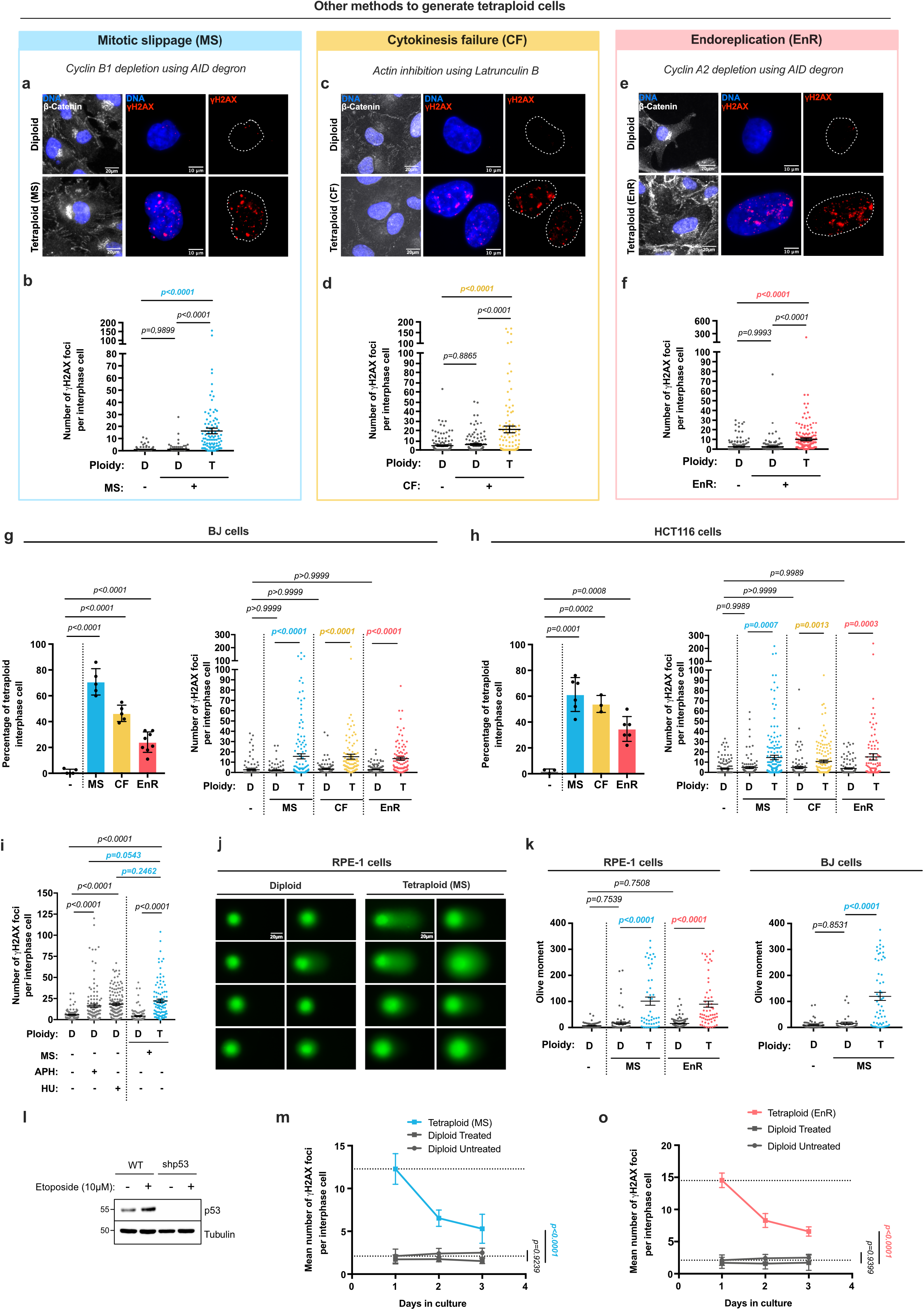
Additional methods and cell lines confirm that WGD generates high levels of DNA damage within the first interphase. **(a, c** and **e)** Images showing diploid and tetraploid (generated as indicated) RPE-1 cells labeled with γH2AX (red) and β-Catenin (gray) antibodies. DNA in blue. **(b, d** and **f)** Graphs showing the number of γH2AX foci in diploid (D) and tetraploid (T) RPE-1. Mean ± SEM, >100 interphase cells were analyzed from at least three independent experiments. **(g-h)** Left - Graph showing the percentage of tetraploid interphase cells in BJ **(g)** or HCT116 **(h)** cell lines. Mean ± SEM, >100 interphase cells were analyzed from at least three independent experiments. Right - Graph representing the number of γH2AX foci in diploid and tetraploid BJ **(g)** or HCT116 **(h)** cells. Mean ± SEM, >100 interphase cells were analyzed from at least three independent experiments. **(i)** Graph showing the number of γH2AX foci in diploid (gray) or tetraploid (blue) RPE-1 cells treated with 1μM APH or 2mM HU. Mean ± SEM, >100 interphase cells were analyzed from at least three independent experiments. **(j)** Comet images from diploid (left) and tetraploid (right) RPE-1 cells. **(k)** Graph showing the olive moment in diploid and tetraploid RPE-1 (left) or BJ (right) cell lines. Mean ± SEM, > 100 comets were analyzed from two independent experiments. **(l)** p53 and tubulin levels assessed by western blot. Etoposide was added as a control for the increased p53 levels. **(m** and **o)** Graphs representing the mean number of γH2AX foci per interphase cell over time (days in culture) in diploid and tetraploid RPE-1 cells. Mean ± SEM, > 100 interphase cells were analyzed from two independent experiments. The dotted lines indicate nuclear area. ANOVA test (one-sided) **(b, d, f, g, h, i, k, m** and **o)**.

**Extended data Figure 3:**
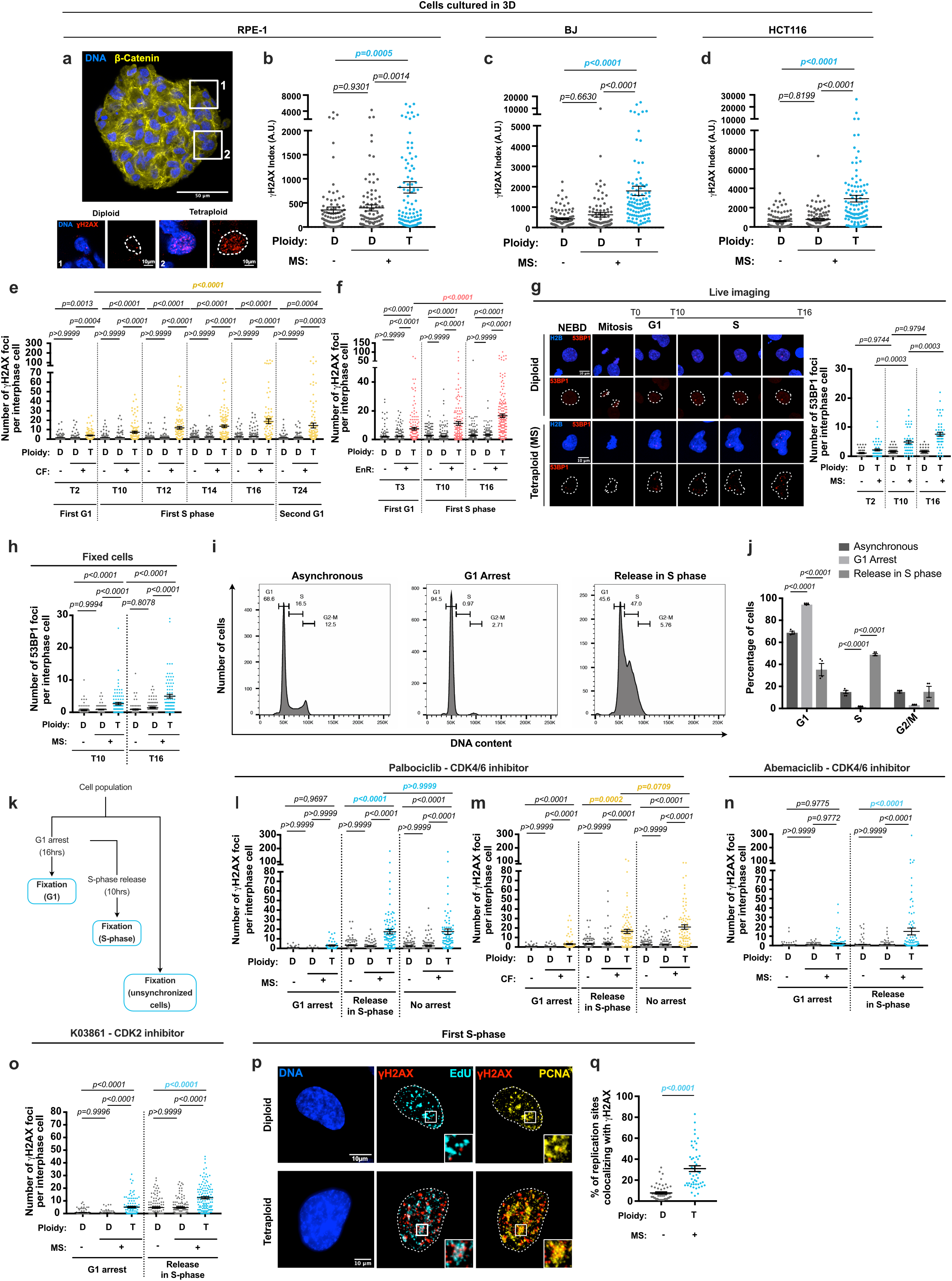
DNA damage is generated during the first S-phase upon WGD. **(a)** 3D RPE-1 spheroid low magnification (top) and insets of two cells showing diploid and tetraploid nuclei (bottom) induced through MS labeled with γH2AX (red) and β-Catenin (yellow) antibodies. DNA in blue. **(b-d)** γH2AX index in diploid and tetraploid RPE-1 **(b)**, BJ **(c)** and HCT116 **(d)** spheroids. Mean ± SEM, > 95 interphase cells were analyzed from at least two independent experiments. (**e-f**) γH2AX foci in diploid and tetraploid RPE-1 cells over time. **(g)** Left - Stills of RPE-1 cells expressing RFP1145-H2B and GFP-53BP1 time lapse videos. Right-53BP1 foci number in diploid and tetraploid cells. Mean ± SEM, > 40 interphase cells were analyzed from three independent experiments. **(h)** 53BP1 foci number in diploid and tetraploid RPE-1. **(i)** Cell cycle distribution of RPE-1 cells in the indicated conditions. **(j)** Percentage of RPE-1 cells in G1, S and G2-M in the indicated conditions. Mean ± SEM, >30 000 cells from at least three independent experiments. **(k)** Workflow used to analyze G1 or S-phase cells. **(l** and **m)** γH2AX foci number in diploid and tetraploid RPE-1 cells as indicated. Experiments **(l** and **m)** share the same reference control. **(n-o)** γH2AX foci number in diploid and tetraploid RPE-1 cells synchronized in G1 using 0,5μM abemaciclib **(n)** or 1μM K03861 **(o)** or released in S-phase. **(p)** Images of γH2AX (red) and EdU (cyan)/ PCNA (yellow) foci co-localization in S-phase in diploid and tetraploid RPE-1 cells. DNA in blue. White squares highlight higher magnifications. **(q)** Percentage of replication sites (EdU) colocalizing with γH2AX foci. Mean ± SEM, >50 interphase cells were analyzed from at least three independent experiments. For (**e**, **f**, **h**, **l**-**o)** Mean ± SEM, >100 interphase cells were analyzed from at least three independent experiments. The dotted lines indicate nuclear area. ANOVA test (one-sided) **(b, c, d, e, f, g, h, j, l, m, n** and **o)**. t-test (two-sided) **(q)**.

**Extended data Figure 4:**
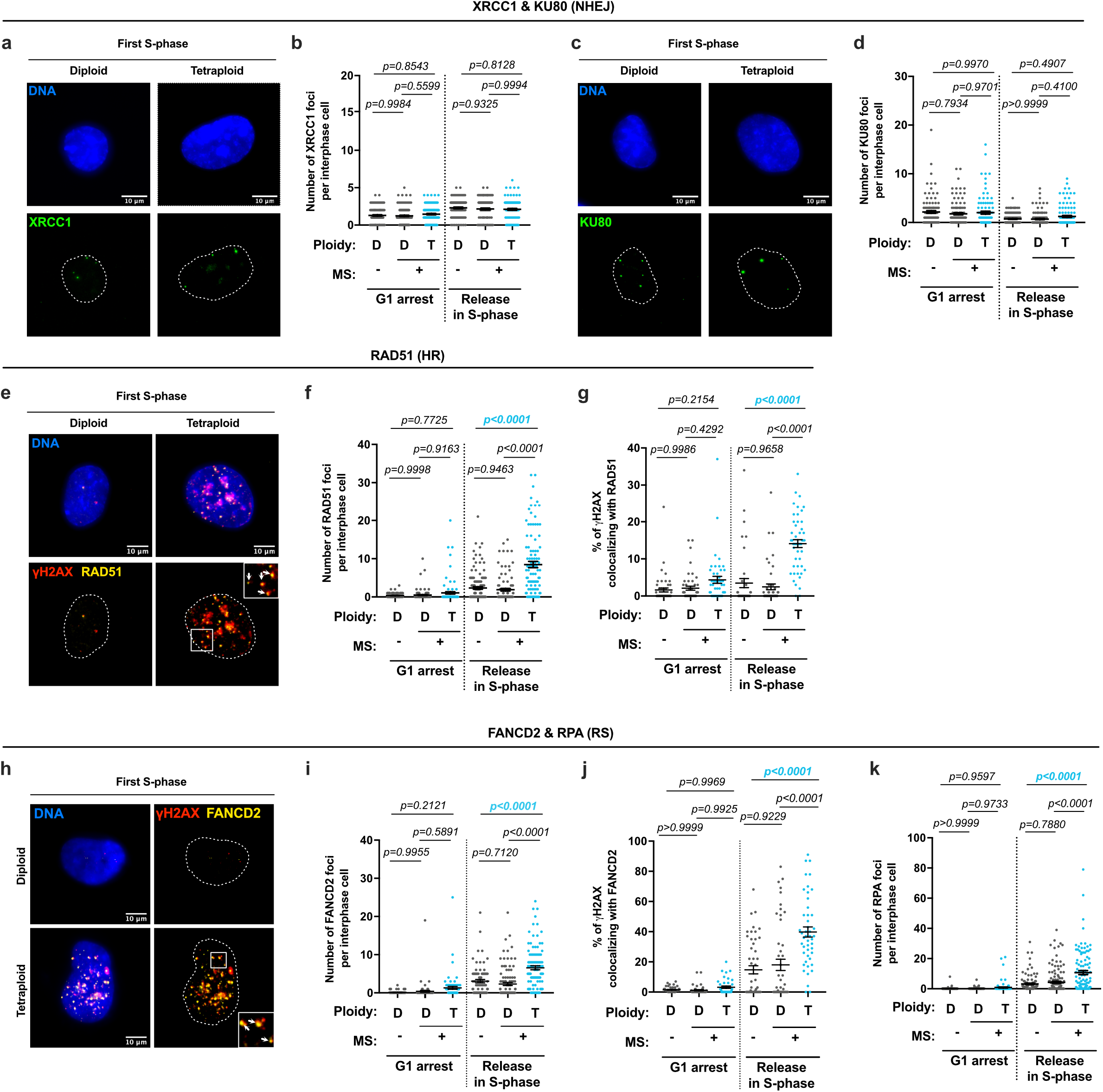
DNA damage in newly born tetraploid S-phase cells is associated with HR and RS-associated markers. **(a** and **c)** Diploid and tetraploid RPE-1 S-phase cells labeled with XRCC1 (**a**, in green) or KU80 antibodies (**c**, in green). DNA in blue. **(b** and **d)** Number of XRCC1 **(b)** or KU80 **(d)** foci in diploid and tetraploid RPE-1 cells synchronized in G1 using 1μM palbociclib or released in S-phase. Mean ± SEM, >100 interphase cells were analyzed from at least three independent experiments. **(e)** Images showing γH2AX (red) and RAD51 (yellow) foci colocalization (white arrows) in diploid and tetraploid RPE-1 cells. DNA in blue. The white squares correspond to higher magnification regions. **(f)** RAD51 foci number in diploid and tetraploid RPE-1 cells arrested in G1 using 1μM palbociclib or released in S-phase. Mean ± SEM, >100 interphase cells analyzed from at least three independent experiments. **(g)** Percentage of colocalizing γH2AX and RAD51 signals in diploid and tetraploid RPE-1 cells arrested in G1 using 1μM palbociclib or released in S-phase. Mean ± SEM, >50 interphase cells were analyzed from at least three independent experiments. **(h)** Images showing the colocalization (white arrows) of γH2AX (red) and FANCD2 (yellow). DNA in blue. **(i)** FANCD2 foci number in diploid and tetraploid RPE-1 cells. Mean ± SEM, >80 interphase cells were analyzed from at least three independent experiments. **(j)** Graph representing the percentage of γH2AX signal colocalizing with FANCD2 foci in diploid and tetraploid RPE-1 interphase cells. Mean ± SEM, >50 interphase cells were analyzed from at least three independent experiments**. (k)** Graph showing RPA number foci in diploid and tetraploid RPE-1 cells arrested in G1 using 1μM palbociclib or released in S-phase. Mean ± SEM, >100 interphase cells were analyzed from at least three independent experiments. The dotted lines indicate the nuclear area. ANOVA test (two sided) **(b, d, f, g, i, j** and **k)**.

**Extended data Figure 5:**
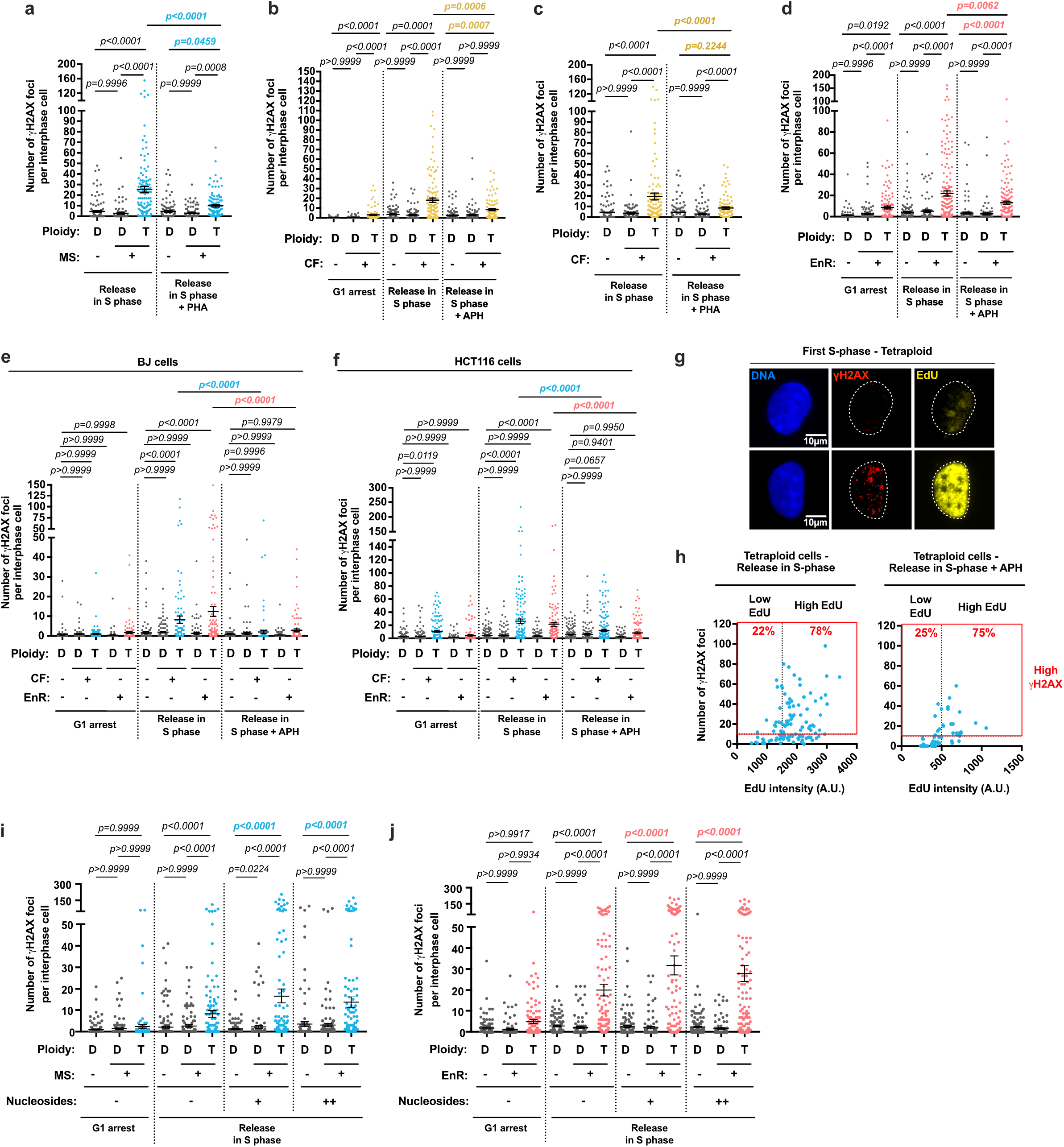
DNA damage in newly born tetraploid cells is generated in a DNA replication-dependent manner. **(a)** γH2AX foci number in diploid and tetraploid RPE-1 cells released in S-phase ± 1μM PHA. Mean ± SEM, >100 interphase cells were analyzed from at least three independent experiments. **(b-c)** γH2AX foci number in diploid and tetraploid RPE-1 cells, arrested in G1 using 1μM palbociclib or released in S-phase ± 400nM aphidicolin (APH) **(b)** or 1μM PHA **(c)**. Mean ± SEM, >100 interphase cells were analyzed from at least three independent experiments. **(d)** γH2AX foci number in diploid and tetraploid RPE-1 cells released in S-phase ± 400nM APH. Mean ± SEM, >100 interphase cells were analyzed from at least three independent experiments. **(e-f)** γH2AX foci number in diploid and tetraploid BJ **(e)** or HCT116 **(f)** cells, released in S-phase ± 400nM APH. Mean ± SEM, >100 interphase cells were analyzed from at least three independent experiments. **(g)** Images showing EdU ± tetraploid RPE-1 cells. γH2AX antibodies in red, EdU in yellow and DNA in blue. **(h)** γH2AX foci number relative to EdU intensity in RPE-1 tetraploid cells released in S1204-phase untreated (left panel) or treated (right panel) with 400nM aphidicolin (APH). Mean ± SEM, >100 interphase cells were analyzed from at least three independent experiments. **(i** and **j)** γH2AX foci number in diploid and tetraploid cells (**i**, blue) or EnR (**j**, red), synchronized in G1 using 1μM palbociclib or released in S-phase ±nucleosides at two different concentrations (methods). Mean ± SEM, >100 interphase cells were analyzed from at least three independent experiments. The dotted lines indicate the nuclear area. ANOVA test (one-sided) **(a, b, c, d, e, f, i** and **j)**. Pearson test (two-sided) **(h)**.

**Extended data Figure 6:**
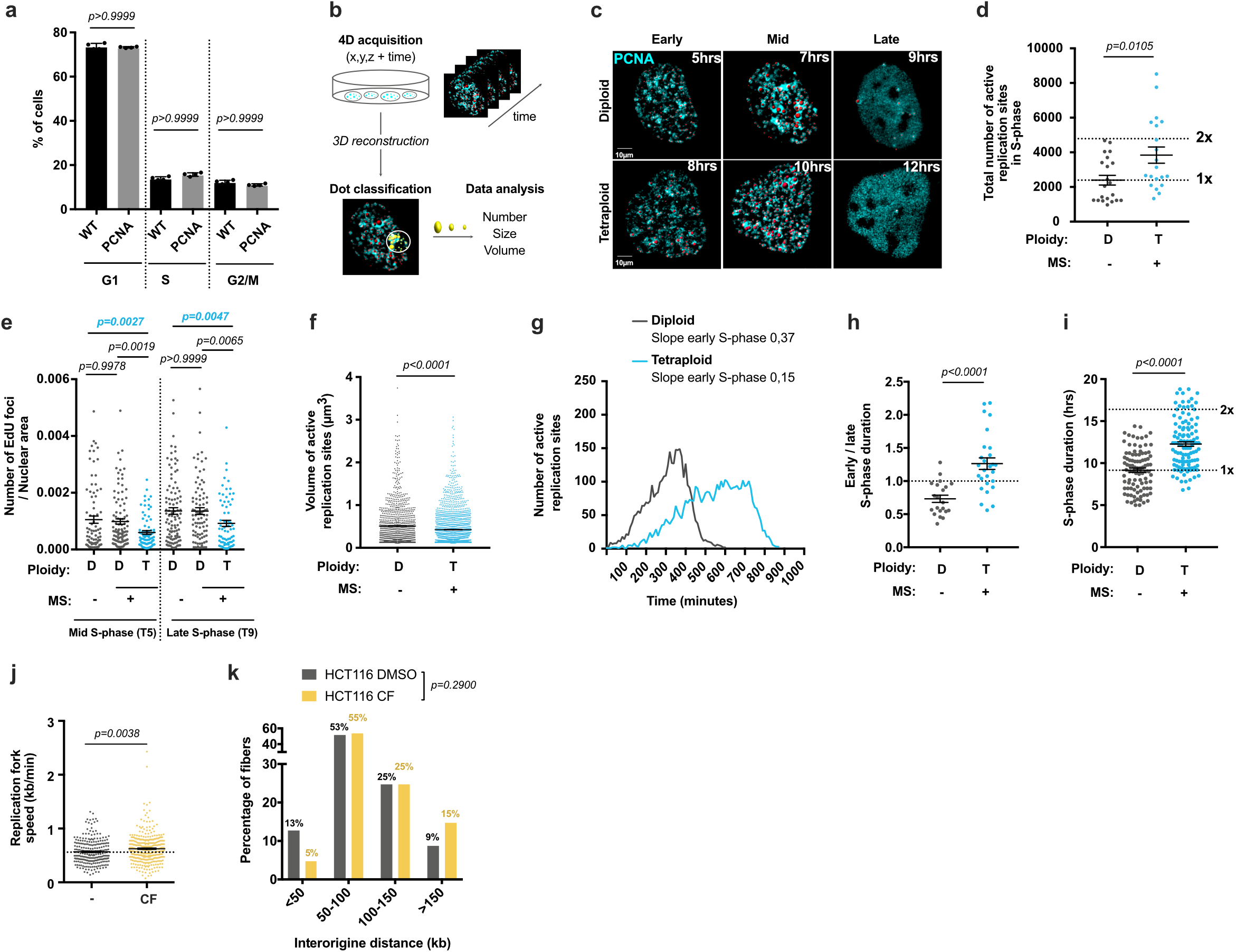
DNA replication dynamics is impaired during the first S-phase in tetraploid cells. **(a)** Percentage of cells per cell cycle phase in RPE-1 (dark gray) and RPE-1 PCNA^chromo^ cell lines (light gray). **(b)** Workflow depicting methods used to process and analyze DNA replication by time-lapse. **(c)** Stills of time lapse movies of diploid and tetraploid RPE-1 PCNA^chromo^ cells. Active replication sites are visualized using PCNA chromobodies (in cyan) and reconstructed using *Imaris* in 3D (in red). **(d)** Total number of active replication sites in S-phase in diploid and tetraploid RPE-1 cells. Mean ± SEM, >20 S-phase cells were analyzed from three independent experiments. **(e)** EdU foci number relative to nuclear area in diploid and tetraploid RPE-1 cells in mid (T5) or late (T9) S-phase. Mean ± SEM, >100 interphase cells were analyzed from at least three independent experiments. **(f)** Volume of active replication sites (in μm3) for diploid and tetraploid RPE-1 PCNA^chromo^ cells. Mean ± SEM, at least 1000 active replication sites were analyzed from three independent experiments. **(g)** Mean number of active replication sites over time in diploid and tetraploid RPE-1 cells. >20 S-phase cells were analyzed from two independent experiments (see Supplemental data 1). **(h)** Ratio of early/late S-phase duration in diploid or tetraploid RPE-1 PCNA^chromo^ cells ± extended G1 duration. Mean ± SEM, > 70 cells from two independent experiments were analyzed. **(i)** S-phase duration in diploid or tetraploid RPE-1 PCNA^chromo^ cells ± extended G1 duration. Mean ± SEM, > 70 cells from two independent experiments were analyzed. **(j)** Replication fork speed in diploid and tetraploid HCT116 cells. Mean ± SEM, > 250 replication forks were analyzed. **(k)** Proportion of fibers with the indicated inter-origin distance (kb) in diploid or tetraploid HCT116 cells. Mean ± SEM, > 75 replication origins were analyzed. ANOVA test (one-sided) **(a** and **e)**. t-test (two-sided) **(d, f, h, i, j** and **k)**.

**Extended data Figure 7:**
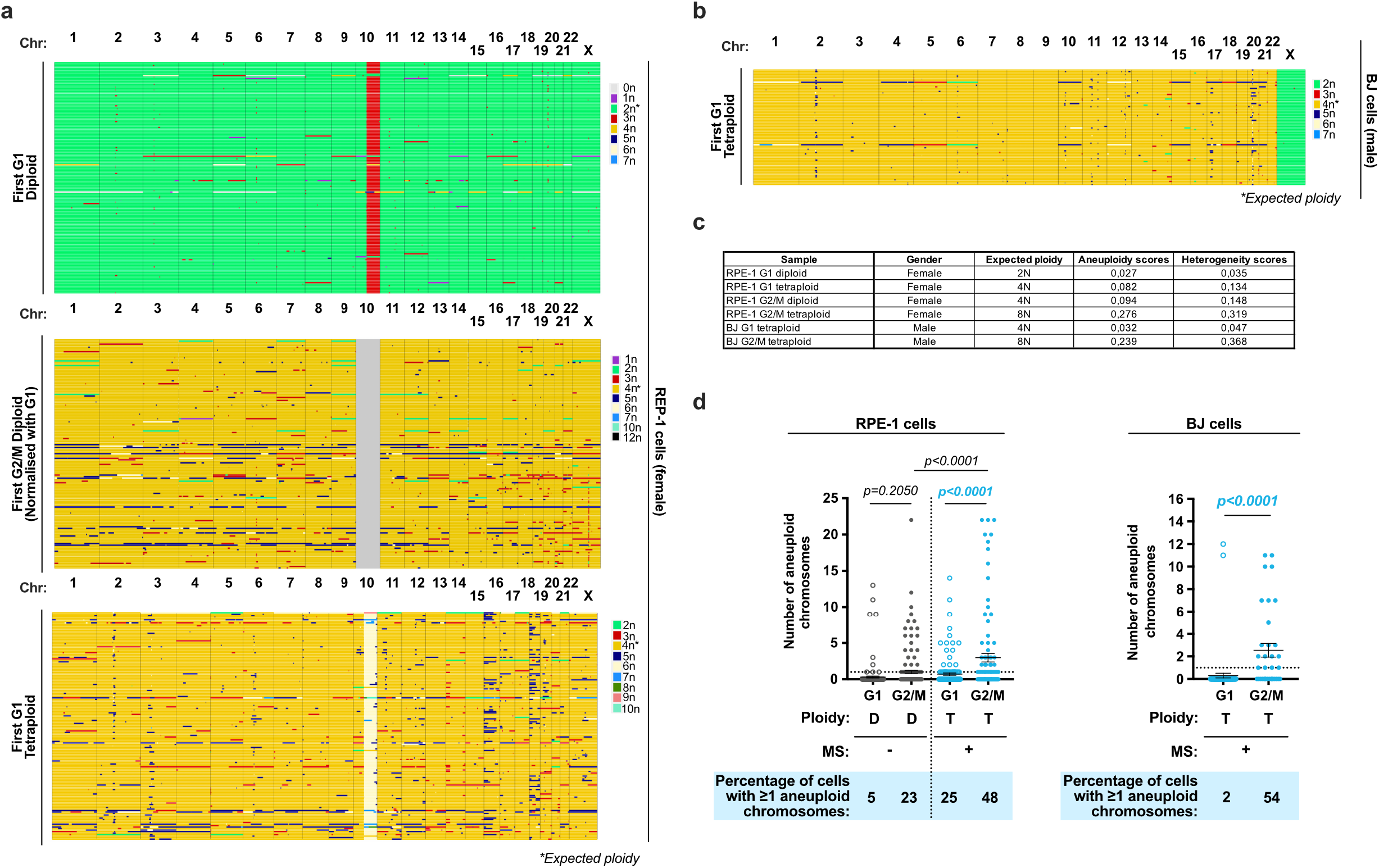
Genome wide analysis of RPE-1 and BJ tetraploid cells. **(a** and **b)** Genome-wide copy number plots of G1 and G2/M diploid RPE-1 cells and G1 tetraploid RPE-1 cells **(a)** were generated using the standard version of the Aneufinder algorithm and genome-wide copy number plots of G1 tetraploid BJ cells **(b)** were generated using a modified version of the Aneufinder algorithm (see methods). G2/M conditions were normalized using G1 cells. Each row represents a cell. The copy number state (in 1-Mb bins) is indicated in color (with aberrations contrasting from green in diploid G1 (2n) or from yellow in diploid G2/M or tetraploid G1 (4n). **(c)** Table showing aneuploidy and heterogeneity scores in the indicated conditions. **(d)** Graph showing the number of aneuploid chromosomes per cell in the diploid G1 and G2/M (in gray) and in tetraploid G1 and G2/M (in blue) cells. The percentage of cells with ≥1 aneuploid chromosome is indicated under the graph. ANOVA test (one-sided) (**d,** left panel). t-test (two-sided) (**d,** right panel).

**Extended data Figure 8:**
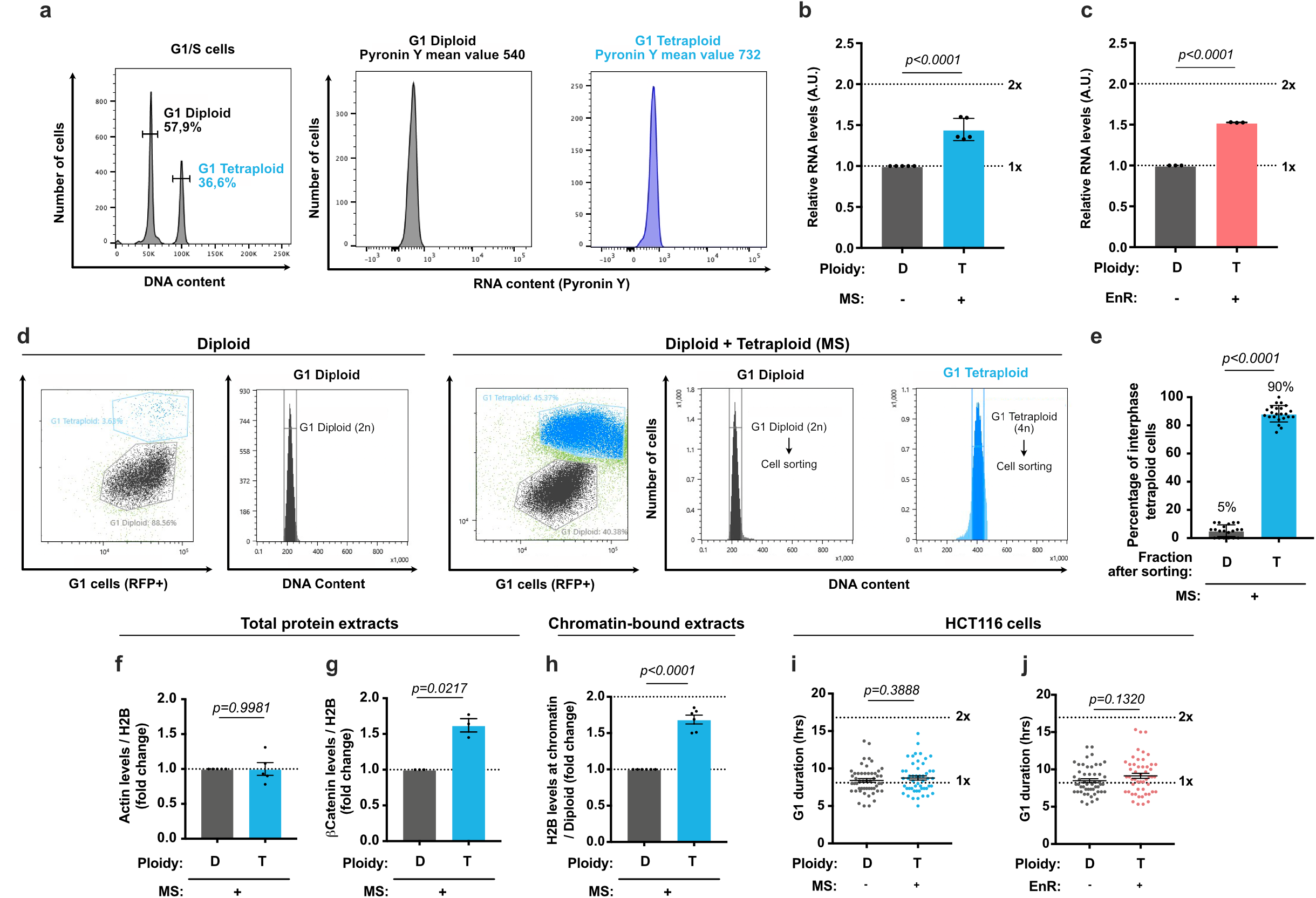
DNA damage analysis in 3D cultures. **(a)** Left panel - Representative cell cycle distribution of diploid and tetraploid RPE-1 cells. Right panels - RNA content in diploid (in gray) and tetraploid (in blue) populations. **(b** and **c)** Graphs showing the relative RNA levels in diploid (D, in gray) and tetraploid (T) cells generated through MS (**b**, blue) or EnR (**c**, red). **(d)** Representative images of cell sorting experiments according to cell cycle stage (RFP+ for G1 cells) and DNA content. **(e)** Graph showing the percentage of interphase tetraploid cells in diploid (gray) and tetraploid (blue) RPE-1 cell populations obtained after cell sorting. Mean ± SEM, > 100 interphase cells from at least three independent experiments were analyzed. **(f** and **g)** Graphs representing actin **(f)** and β-Catenin **(g)** levels relative to H2B levels (fold change) in total protein extracts from diploid (gray) and tetraploid (blue) cells. Mean ± SEM from at least three independent experiments. **(h)** Graph showing H2B levels in the chromatin bound fraction in diploid (gray) and tetraploid (blue) cells. Mean ± SEM from at least three independent experiments. **(i** and **j)** Graph representing G1 duration in diploid (gray) or tetraploid cells generated through MS (**i**, blue) or EnR (**j**, red). The dotted lines indicate the nuclear area. The white squares correspond to higher magnification. t-test (one-sided) **(b, c, e, f, g, h, i** and **j)**.

**Extended data Figure 9:**
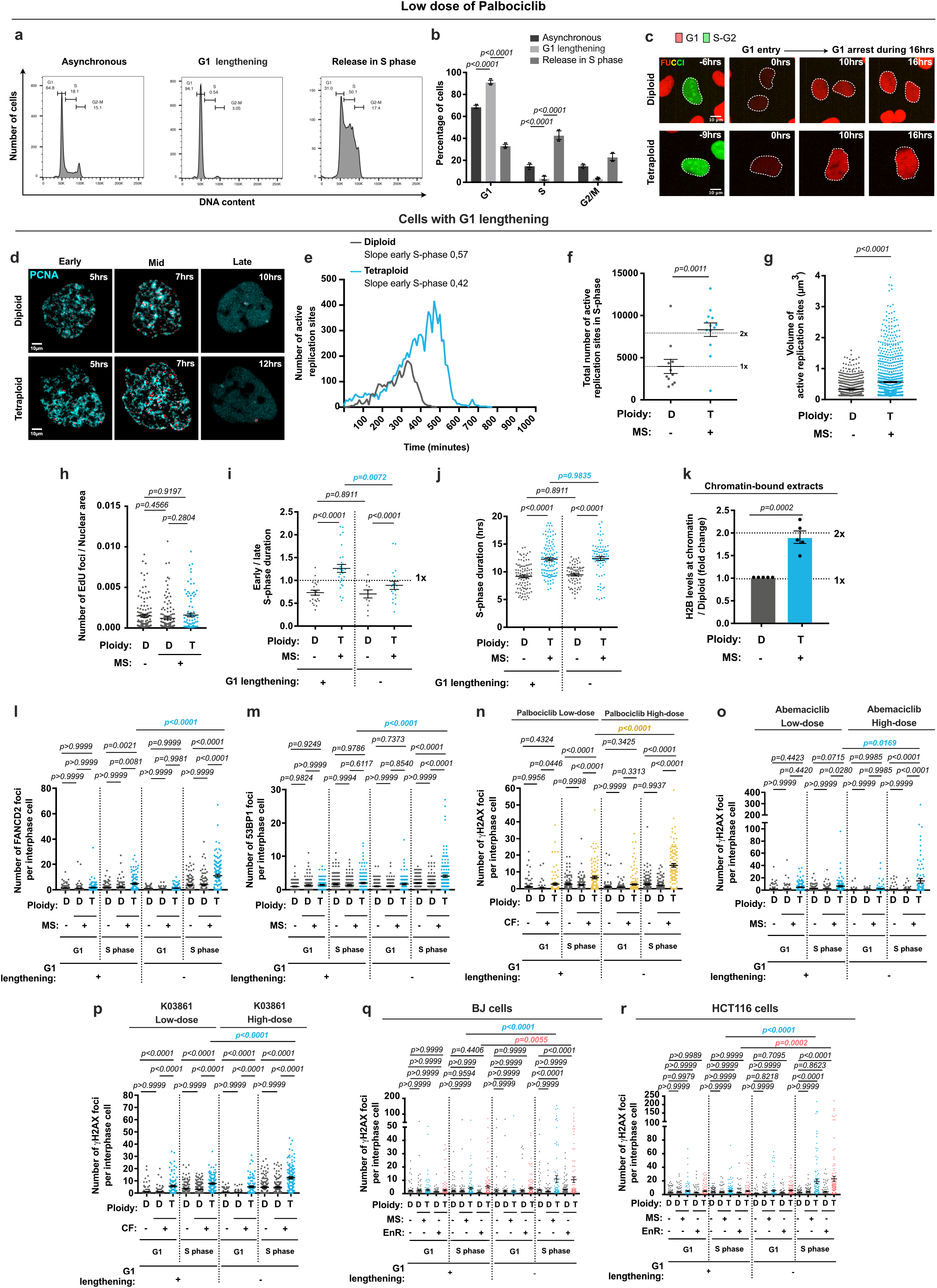
G1 lengthening restores DNA replication dynamics and results in a decrease in the levels of DNA damage in tetraploid cells. **(a-b)** RPE-1 cell cycle profile and percentage of cells in the indicated conditions. Mean ± SEM, > 30 000 cells from at least three independent experiments. **(c)** RPE FUCCI in diploid and tetraploid cells treated with 160nM palbociclib. **(d)** Stills of time lapse videos of diploid and tetraploid RPE-1 PCNA^chromo^ cells with extended G1. Active replication sites visualized using PCNA chromobodies (cyan) and reconstructed using *Imaris* in 3D (red). **(e)** Active replication sites average number over time with extended G1. Mean ± SEM, > 11 S-phase cells analyzed, two independent experiments (see Supplemental data 1). **(f)** Active replication sites total number with extended G1. Mean ± SEM, > 11 S-phase cells were analyzed, two independent experiments. **(g)** Active replication sites volume (μm3) with extended G1. Mean ± SEM, > 1000 Active replication sites analyzed, three independent experiments. **(h)** EdU foci number relative to nuclear area with extended G1. Mean ± SEM, > 100 interphase cells, at least three independent experiments. **(i)** Ratio of early/late S phase duration ± extended G1. Mean ± SEM, > 70 cells, two independent experiments. **(j)** S-phase duration ± extended G1. Mean ± SEM, > 70 cells, two independent experiments. **(k)** H2B levels in chromatin bound extracts. Mean ± SEM, four independent experiments. **(l** and **m)** FANCD2 or 53BP1 foci number in cells synchronized in G1 or released in S-phase ± extended G1. **(n-p)** γH2AX foci number in cells synchronized in G1 using the indicated treatments or released in S-phase ± extended G1. **(q** and **r)** γH2AX foci number in diploid and tetraploid BJ **(q)** or HCT116 **(r)** cells synchronized in G1 or released in S-phase ± extended G1. **(l-r)** Mean ± SEM, >100 interphase cells were analyzed, at least three independent experiments. The dotted lines indicate the nuclear area. ANOVA test (one-sided) **(b, h, i, j, l, m, n, o, p, q** and **r)**. T-test (two-sided) **(f, g** and **k)**.

**Extended data Figure 10:**
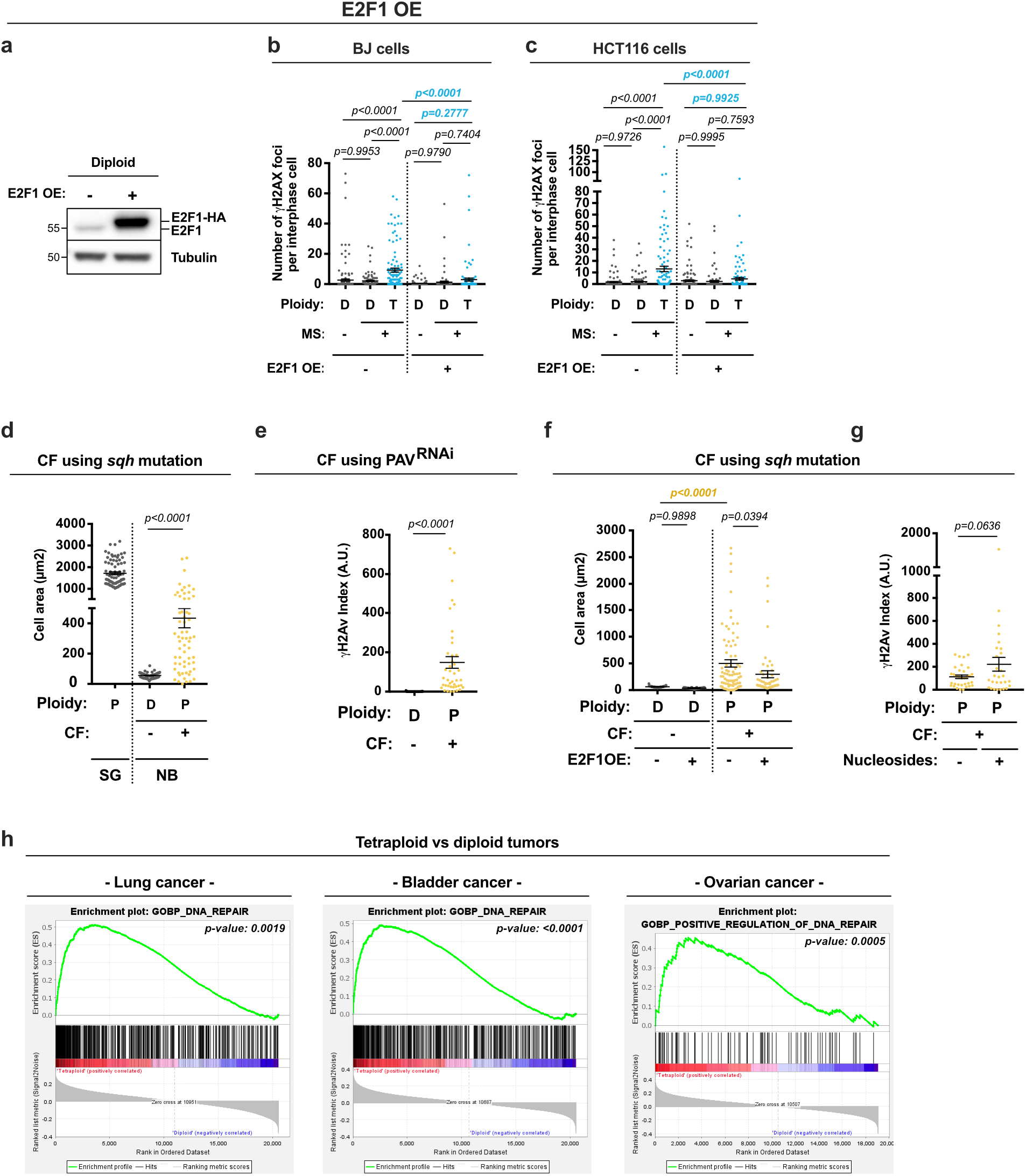
E2F1OE decreases DNA damage levels in tetraploid human cell lines and in *Drosophila* NBs. **(a)** Western blot documenting the levels of E2F1 and tubulin from cell lysates obtained from diploid RPE-1 cells ± E2F1-HA over-expression (OE). **(b-c)** Graphs representing the number of γH2AX foci per interphase cell in diploid (D) and tetraploid (T) BJ **(b)** or HCT116 **(c)** cells released in S-phase ± E2F1 OE. Mean ± SEM, >100 interphase cells were analyzed from at least three independent experiments. **(d)** Graph showing wild type salivary gland cell (in gray), diploid (in gray) and polyploid NBs (in yellow) area (in μm2). Mean ± SEM, >60 cells were analyzed per condition. **(e)** Graph showing γH2Av indexes in diploid (in gray) or polyploid NBs (in yellow) induced through CF by depleting Pavarotti. Mean ± SEM, >40 cells were analyzed per condition. **(f)** Graph showing the cell area (μm^2^) of diploid (gray) and polyploid NBs (yellow) ± E2F1OE. Mean ± SEM, >30 cells were analyzed per condition. **(g)** Graph representing the γH2Av index in polyploid NBs ± 10μM nucleosides. Mean ± SEM, >28 cells were analyzed per condition. **(h)** Gene set enrichment analysis from GSEA. Plots show significant enrichment of DNA repair genes in near-tetraploid tumors when compared to near-diploid tumors in lung, bladder and ovarian cancers (TCGA pan cancer data set). **(h)** p value from false discovery rate (FDR; methods). ANOVA test (one-sided) **(b, c, d, f)**. t-test (two-sided) **(e** and **g)**.

