## Supplementary material for "Genetic instability from a single S-phase after whole genome duplication": SI Guide

7- Present address: Molecular, Cellular, and Developmental Biology Department, University of California, Santa Barbara, CA, USA.

### SUPPLEMENTAL INFORMATION GUIDE


**Supplementary Information** **p.3**

**Supplementary Discussion** **p.5**

**Supplementary References** **p.7**

**Supplementary Methods 1** **p.8**

**Supplementary Data 1** **p.9**

**Supplementary Data 2** **p.10**

**Figure legends for extended videos** **p.12**

**Supplementary Information:**

#### PCNA analysis

To validate the use of RPE-1 cells stably expressing PCNA Chromobodies ($PCNA^{chromo}$ cells), we measured cell cycle progression and compared with control cells. We found that PCNA chromobodies did not affect DNA replication or cell cycle progression (Extended data Fig.6a). Concerning PCNA behavior in diploid cells, as S-phase is initiated, an exponential increase in the number of active replication sites was noticed, which was maintained before undergoing a rapid decrease as cell exited S-phase. In contrast, in tetraploid cells the increase in the number of active sites was more gradual, and the signals associated with DNA lingered for extended periods of time (compare slope curves in diploid- 0.37 and tetraploid- 0.15 cells; Extended Fig. 6g). After G1 extension, the dynamic behavior of PCNA in tetraploid cells was comparable to diploid cells (compare slope curves after extended G1 in diploid- 0,57 and tetraploid- 0,42 cells; Extended data Fig. 9d-h and Extended data movies 5-6).

#### Single Cell sequencing of tetraploid cells and respective controls

With the aim of separating from the same population diploid from tetraploid cells, we developed a protocol that allows Fluorescence Activating Cell Sorting (FACS) based on DAPI intensity and cell cycle markers (Figure 3c and Extended data Fig. 8d-e and methods). Tetraploid and diploid cells were sorted at G1/S and at G2/M transitions. Normalization of the under- and over-replicated regions at G1/S and G2/M diploid cells already revealed whole chromosome deviations in a small number of cells. When present, they span along almost all chromosomes of a given cell (Extended data Fig. 7a-b). In G1 tetraploid cells, over-replicated regions (5n) could also be identified, but these were restricted to a few chromosomes and might be explained by a caveat of the method (cells have initiated S-phase but were still selected as G1 by the FACS profile). In G2/M tetraploid cells, however over-duplicated chromosomes (> 10) were identified in addition to frequent over- and under-replicated regions (9n, 7n and 4n) (Fig. 2h-i).

By analyzing G1 and G2/M tetraploid cells, we concluded that defects in S-phase result in the generation of highly aberrant karyotypes, demonstrating a causal relationship between tetraploidization and GIN within a single S-phase.

**Use of CDK4/CDK6 and CDK2 inhibitors to extend G1**

To induce G1 lengthening, we used low doses of Palbociclib, Abemaciclib and K03861 (methods). These conditions were different from the ones described earlier in this study to synchronize cells in G1. Indeed, while high doses of these inhibitors result in a cell cycle arrest, low inhibitor doses result in G1 lengthening (Extended data Fig. 9a-c) [1,2]. The different impact of high and low doses of CDK4/6 or CDK2 inhibitors could be noticed in differences in the expression levels of DNA replication factors (Figure 3e-h vs Figure 3k-o).

**Supplementary Discussion:**

Analysis of tetraploid cells treated with increasing levels of nucleosides did not rescue high levels of DNA damage in the first interphase typical of these cells (Extended data Fig. 5i-j). This was also the case in experiments performed *in vivo* using the non-physiological polyploid model system- *Drosophila* NBs (Extended data Fig. 10g). Collectively these data suggest that nucleoside addition does not rescue DNA replication defects reported here. At least two plausible explanations may be considered. The first hypothesis is that total nucleoside levels are not a limiting factor in tetraploid cells as it is the case for other factors described here (Figure 3). Therefore, their addition does not ameliorate the capacity of tetraploid cells to undergo S-phase in an optimal manner. The second hypothesis is that even if nucleoside addition may enhance S-phase fidelity, the fact that other key S-phase replication factors are still not scaled up, it is not possible to rescue genetic instability and DNA damage.

One of the most surprising results of this study was the accelerated fork progression noticed in DNA combing approaches in all tetraploid induced cell lines and this even by different means - CF, MS and EnR (Figures 2f-g and Extended data Fig. 6j). Moreover, marked defects in fork symmetry, which are consistent with fork collapse were also noticed in all the conditions (Figure 2f-g). It is important to mention that the increased RAD51 and FANCD2 levels in tetraploid cells suggest a role in restarting stalled replication forks as the DDR is normally initiated [3,4]. To distinguish between possible defects in either pre-RCs loading in G1 and DNA replication initiation factors, normally loaded in S-phase, we tested their levels in chromatin-bound extracts. Importantly, proteins from both categories did not show the expected scaling up factor (Figure 3e and g). Extended G1 was sufficient to rescue protein levels in tetraploid cells and to decrease DNA damage levels (Figure 3k-q). Together, it is possible to conceive at least two likely, not mutually exclusive, scenarios to explain the origins of DNA damage levels described here. The first is that the low levels of Pre-RCs may account for increased fork speed as shown in yeast [5] or more recently upon decreased MCM levels in mammalian cells [6]. The second possibility, concerns the observations that origin firing factors such as Treslin or PCNA and CDC45, members of the active replisome, did not scale up in tetraploid cells. Importantly, Treslin depletion also results in increased fork speed as shown in [7]. Together, these results suggest that tetraploid cells are penalized twice having both low levels of Pre-RCs and active replication factors resulting in increased fork speed and asymmetry leading to fork stalling and ultimately to DNA damage. G1 extension or an increase in the expression levels of replication proteins and their association with the DNA is sufficient to lower considerably DNA damage levels. These results highlight the importance of maintaining the balance between DNA and protein content to ensure genetic stability and cell homeostasis.

The most unexpected finding of this study is the lack of scaling up between DNA and protein content immediately after tetraploidization. In physiological conditions, such as during animal development, WGDs and polyploidization lead to an overall scaling up of cell mass and DNA content to favor increase secretion and metabolic activity, for example [8]. Our work shows that unscheduled tetraploid or polyploid cells do not increase cell mass as expected in the first interphase after WGD. Why certain key cell cycle and DNA replication factors fail to be expressed at levels that allow optimal DNA replication remains to be explained. Importantly, however, these results show that an immediate consequence of unscheduled genome doubling is the loss of genetic integrity within a single S-phase. Interestingly, studies performed on stable tetraploid cells have shown scaling up of protein and DNA content after long-term adaptation [9,10], suggesting that evolved tetraploids develop specific mechanisms to bypass the initial lack of scaling up.

#### Supplementary Methods 1: Methods for sequencing analysis

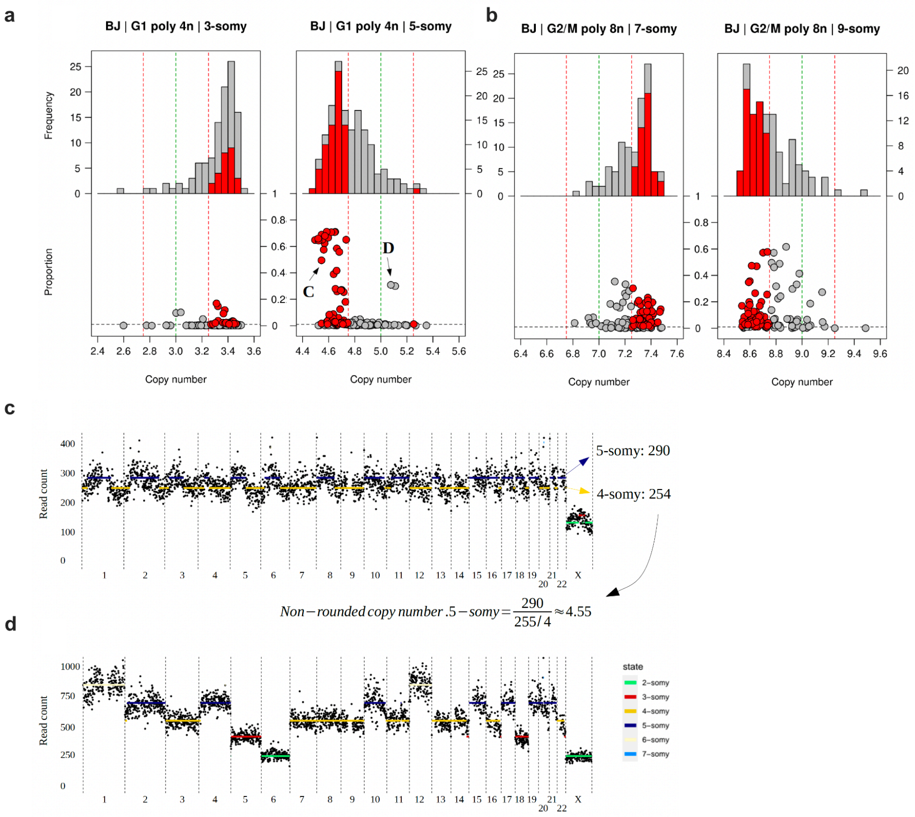


**(a-b)** Distribution of non-rounded copy numbers using either 4-somy (a) or 8-somy (b) as reference. The states that were inspected have either one copy less (3 or 7-somy) or one copy more (5 or 9-somy) compared to the expected state (4 or 8-somy). The green dotted lines indicate the expected value and the red dotted lines indicate the applied cutoffs (deviation of 0.25). Libraries that were removed are indicated with red. The bottom panel shows the proportion of the genome which has the state. The horizontal black dotted line indicates the 1 % cutoff that was applied. The non-rounded copy number of 5-somy (**a**, 4n sample) show for some libraries a tendency towards 4-somy (lower non-rounded values). **(c)** Example of an aberrant library. The means of 4 and 5-somy are too close to each other. **(d)** Example of a clear aneuploid library.

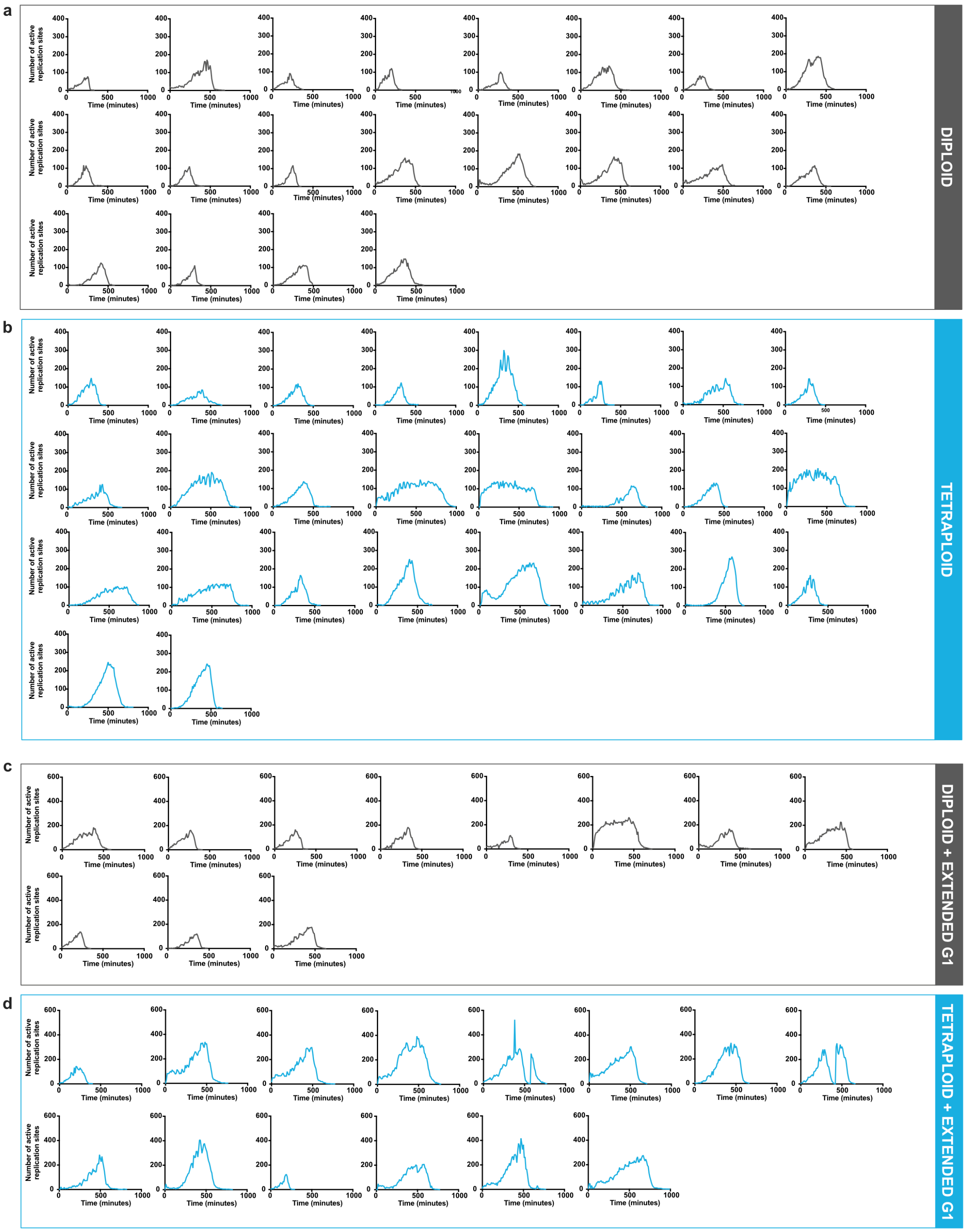


**Supplementary Data 1: Analysis of DNA replication dynamics in RPE-1 cells**
**(a-b)** Profiles showing the average number of active replication sites over time in diploid (**a**, gray lines) or tetraploid (**b,** blue lines) RPE PCNA$^{chromo}$ cells. **(c-d)** Profiles showing the average number of active replication sites over time in diploid (**c**, gray line) or tetraploid (**d**, blue line) RPE PCNA$^{chromo}$ cells with extended G1.

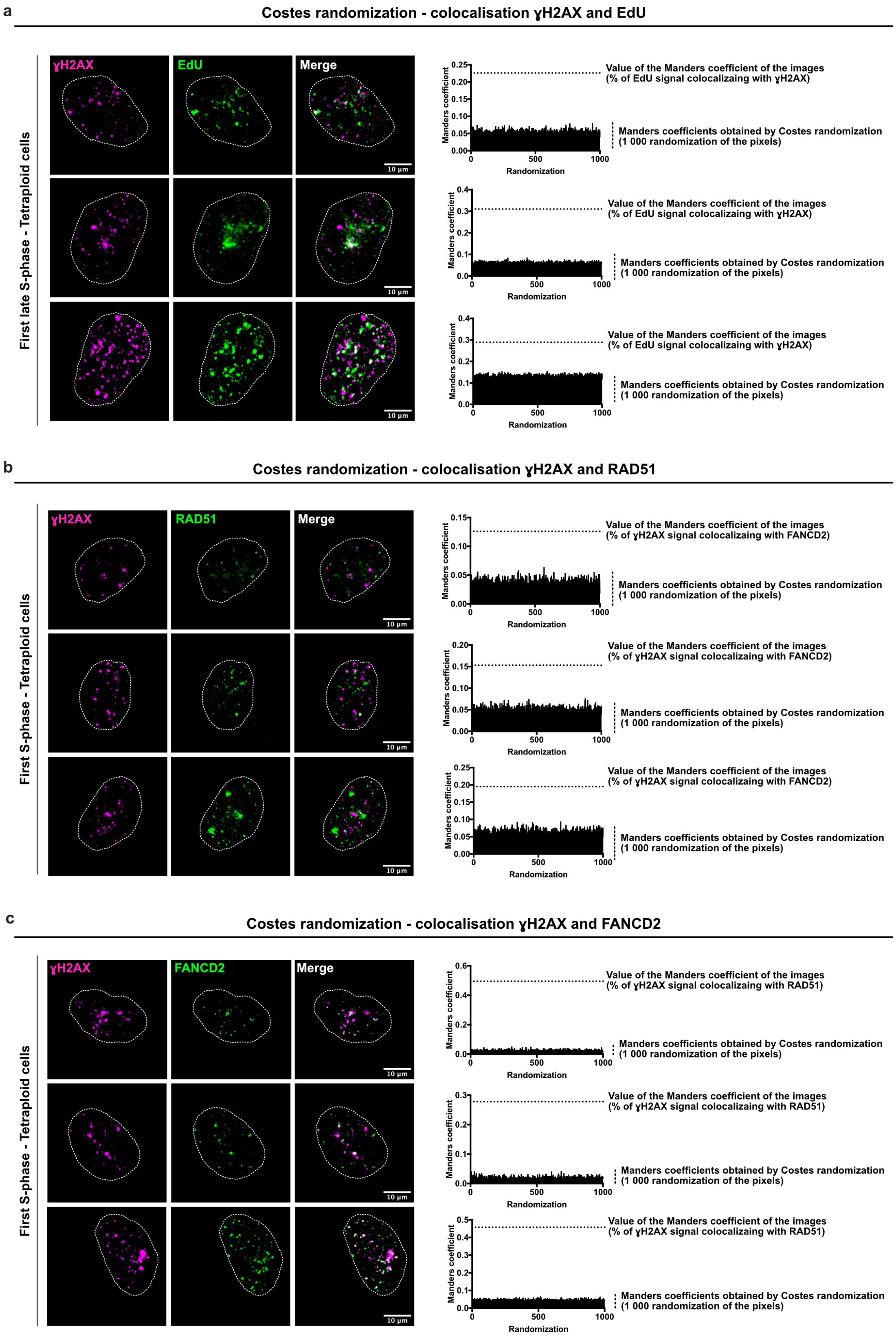


**Supplementary Data 2: Costes randomization for the colocalization of ɣH2AX with EdU / RAD51 or FANCD2.**

**(a-c)** Left panels – Images of S-phase tetraploid cells labeled with antibodies against ɣH2AX (in magenta), EdU (**a**, green), RAD51(**b**, green) and FANCD2 (**c**, green). The dotted lines indicate the nuclear area. Right panels – Results of the home made Costes randomization using the plugin JACOP with Fiji showing that colocalization between ɣH2AX and EdU, RAD51 or FANCD2 cannot result from chance. The analysis was done based on images obtained from 3 independent experiments (n=1000 randomization). The dotted lines indicate the values of Manders coefficient obtained from pictures shown on the left. The Manders coefficient obtained after randomization of the pixels are represented in the graph.

#### FIGURE LEGENDS FOR EXTENDED VIDEOS:

**Extended Videos 1 and 2: DNA damage in tetraploid is generated during S-phase**
Time lapse Videos of RPE-1 diploid (video 1) and tetraploid (video 2) cells expressing RFP-H2B (in blue) and GFP-53BP1 (in red). Images were acquired every 15 minutes.

**Extended Videos 3-6: Quantitative 4D live imaging of endogenous DNA replication.**
Time lapse Videos of RPE-1 diploid (video 3) or tetraploid (video 4) S-phase cells expressing PCNA chromobodies (in cyan). Time lapse Videos of RPE-1 diploid (video 5) or tetraploid (video 6) S-phase cells expressing PCNA chromobodies (in cyan) and treated with 160nM palbociclib before S-phase entry. Images were acquired every 15 minutes.
